# Acquisition of phthiocerol dimycocerosates enabled extension of fused mycomembranes for cellular cording in *Mycobacterium tuberculosis*

**DOI:** 10.64898/2026.11.24.769003

**Authors:** Kenny Jungfer, Lalita Ramakrishnan, Tanmay A. M. Bharat

## Abstract

*Mycobacterium tuberculosis* and its pathogenic relative *Mycobacterium marinum* form virulence-associated serpentine cords. Using live imaging and cryo-electron tomography, we show that cording involves extensive outer membrane enmeshment, such that adjacent cells share a single outer membrane. Importantly, analysis of diverse non-cording *Mycobacteriales* species, including the poorly virulent saprophyte *M. smegmatis,* reveals some membrane enmeshment in the clumped colonies. Our analyses suggest that cording was acquired in two evolutionary steps. First, acquisition of the mycolic acid–rich envelope in *Mycobacteriales* enabled daughter cell membranes to fuse during division, generating membrane-connected clonal clumps. Subsequent acquisition of the virulence-associated outer membrane lipids phthiocerol dimycocerosates (PDIM) facilitated more extensive membrane fusion into cords. We show that PDIM moves freely through the continuous shared outer membranes of cords. Moreover, these *in vitro* observations of fused membranes in *M. marinum* were seen during zebrafish infection. Thus, mycobacterial cording relies on an ancestral fusion-based community architecture that PDIM refined into virulence-promoting cords.

**Significance Statement:** Since its discovery, *Mycobacterium tuberculosis* has been recognized to form cord-shaped cellular assemblies, a property that has been linked to pathogenesis. Using a series of microscopic and microbiological experiments, we show how mycobacterial cord-shaped cellular assemblies are formed by fusing of the outer membranes of bacteria. Analysis of mutants, diverse non-cording mycobacteria and related species identifies the molecular factors important for outer membrane fusion, leading to an evolutionary model of how cording arose in *M. tuberculosis* and its close pathogenic relative *Mycobacterium marinum*.

## Introduction

*Mycobacterium tuberculosis* assumes a distinctive corded appearance during growth in culture that is characterized by elongated, intertwined serpentine rope-like structures (1–3). Cording can be used as a rapid, presumptive diagnostic feature to distinguish the *M. tuberculosis* complex from nontuberculous mycobacteria (4). In animal models, *M. tuberculosis* cords are observed in vivo in granulomas, that are hallmark macrophage-dense immunopathological structures in which the bacteria reside during infection (5–7). Although reports of corded *M. tuberculosis* in human granulomas are scarce, they are seen in the sputum of individuals with TB (8). Mycobacterial cording has long been linked to virulence (9) and is also proposed to act as a physical barrier to antibiotics, compounding the intrinsic antibiotic resistance conferred by the complex mycobacterial cell envelope (7, 10, 11). How cords form, and what surface changes are associated with the transition from single cell to corded growth remain central questions in mycobacterial pathogenesis.

Here, we used live imaging and electron cryotomography (cryo-ET) to examine the architecture of *M. tuberculosis* cords and those formed by its close pathogenic relative, *M. marinum,* where cording is a feature of progressive infection in the zebrafish-tuberculosis model (12, 13). We show that cords grow clonally, as daughter cells stay connected after division via a continuous enmeshed outer membrane (mycomembrane) cell-cell junction. We also find these junctions in non-cording *Mycobacteriales*, including mycobacterial species that form disordered clumps. We further show that phthiocerol dimycocerosates (PDIM), virulence-associated mycomembrane lipids (14–18) are required for the extension of these enmeshed cell-cell junctions and bacteria lacking surface PDIM expression form disorganized clumps rather than cords. Thus, the mycomembrane possesses an intrinsic, ancestral mechanism for linking daughter cells into clonal communities through mycomembrane enmeshment. In the pathogenic species *M. tuberculosis* and *M. marinum*, PDIM enables the extension of these division-generated membrane connections to organize cells into cords.

## Results

### *M. tuberculosis* cords have shared mycomembrane interfaces between apposing cells

To determine their ultrastructure, we used cryo-ET to examine *M. tuberculosis* cords formed during axenic growth in liquid culture (Figure 1A and S1A-D). Canonical Gram-positive and Gram-negative multicellular bacterial communities (biofilms) are typically comprised of discrete individual cells embedded in an extracellular matrix of filamentous proteins, DNA and polysaccharide (19–22). In contrast, we found no visible extracellular material in and around *M. tuberculosis* cords; rather they comprised tightly apposed cells with no gaps between their cell envelopes at points of contact (Figure 1B-F; S2A). To further assess the topology of the interface, we performed subtomogram analysis (23) of the cell-cell contact regions, comparing these with free cell surfaces away from the contact sites (Figure 1G and H).

**Figure 1.**
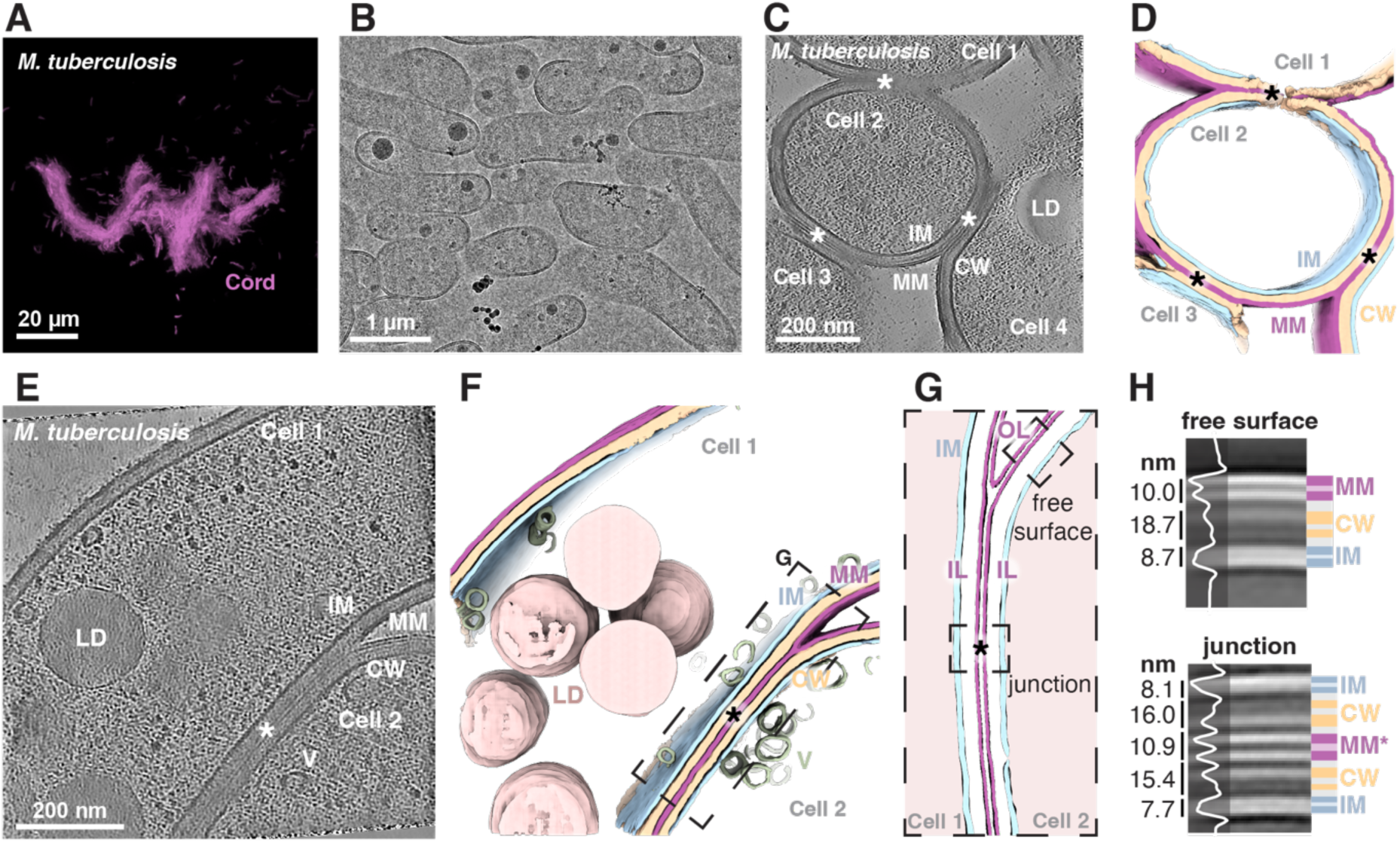
Cells within mycobacterial cords are joined by a single, shared mycomembrane. (A) Fluorescence microscopy of *M. tuberculosis* cords. Cells are depicted in magenta. (B-H) Cryo-ET of *M. tuberculosis* cords. (B) Low magnification overview image of a FIB milled cord. (C) Tomographic slices through cord cell-cell contacts. (D) Three-dimensional segmentation of the same tomogram shown in panel C, showing inner membrane (IM, blue), cell wall and periplasm (CW, yellow), mycomembrane (MM, magenta) and enmeshed mycomembrane contacts (asterisks). (E) Tomographic slice and (F) corresponding segmentation of close cell-cell contact. Cytoplasmic lipid droplets (LD) and vesicles (V) have been highlighted in the segmentation. (G) Segmentations of the mycomembrane junction between adjoining cells showing the inner leaflets (IL) and outer leaflets (OL) of the mycomembrane. (H) Subtomogram analyses and density profiles of the entire cell envelope across the free surface and at the cell-cell junction, with different layers marked.

As expected, the free cell surface of the *M. tuberculosis* envelope showed the expected layers of the mycobacterial envelope - the inner membrane, cell wall and outer membrane (mycomembrane), a highly asymmetric bilayer, containing an inner leaflet formed by mycolic acids covalently linked to the arabinogalactan-peptidoglycan cell wall and an outer leaflet comprising free lipids (24) (Figure 1H). At the interface between apposed cells, the inner membrane and cell wall from each cell were visible; however, instead of four mycomembrane leaflets (two from each cell), only two leaflets were visible (Fig 1H). This topology was consistent with a single bilayered interbacterial membrane shared by the two cells. This was supported by the finding that inter-leaflet distances at contact sites and free surfaces were similar (10.2 nm versus 9.1 nm) (Figure 1H). Following the mycomembrane from the free surface into the contact, we saw that the inner leaflets of the two apposed cells continued into the interface, where they came together to form a single bilayer (Figure 1G). The outer leaflets were absent from the interface. Instead, they curved at the junction and ran continuously around the outside of both cells, enclosing them within one uninterrupted outer leaflet (Figure 1G).

Thus, *M. tuberculosis* community architecture appears to be facilitated not by separate cells connected by extracellular material nor by intact mycomembranes tightly pressed against each other (20). Rather, cords are characterized by direct cell envelope continuity in apposed cells at areas of cell-cell contact. The enmeshed mycomembranes are formed by mycomembrane outer leaflets wrapping around adjacent cells and a new shared bilayer comprising their inner mycomembrane leaflets.

### Mycobacterial cell-cell contacts in vivo are characterized by mycomembrane enmeshments

*M. marinum*, a close pathogenic relative of *M. tuberculosis*, also forms cords, which are morphologically similar to those of *M. tuberculosis* both during axenic and extracellular growth in necrotic granuloma cores (12, 13) (Figures 2A, compare to Figure 1A). Cryo-ET of *M. marinum* cords formed during axenic culture (Figure S1E) confirmed that cell-cell contacts were also characterized by the enmeshment of mycomembranes and a lack of extracellular matrix, similar to those of *M. tuberculosis* cords (Figure 2B-D, Figure S2B). These enmeshed contacts were extensive, with a single continuous membrane spanning multiple cell-cell interfaces (Figure 2B-D, Figure S2B).

**Figure 2.**
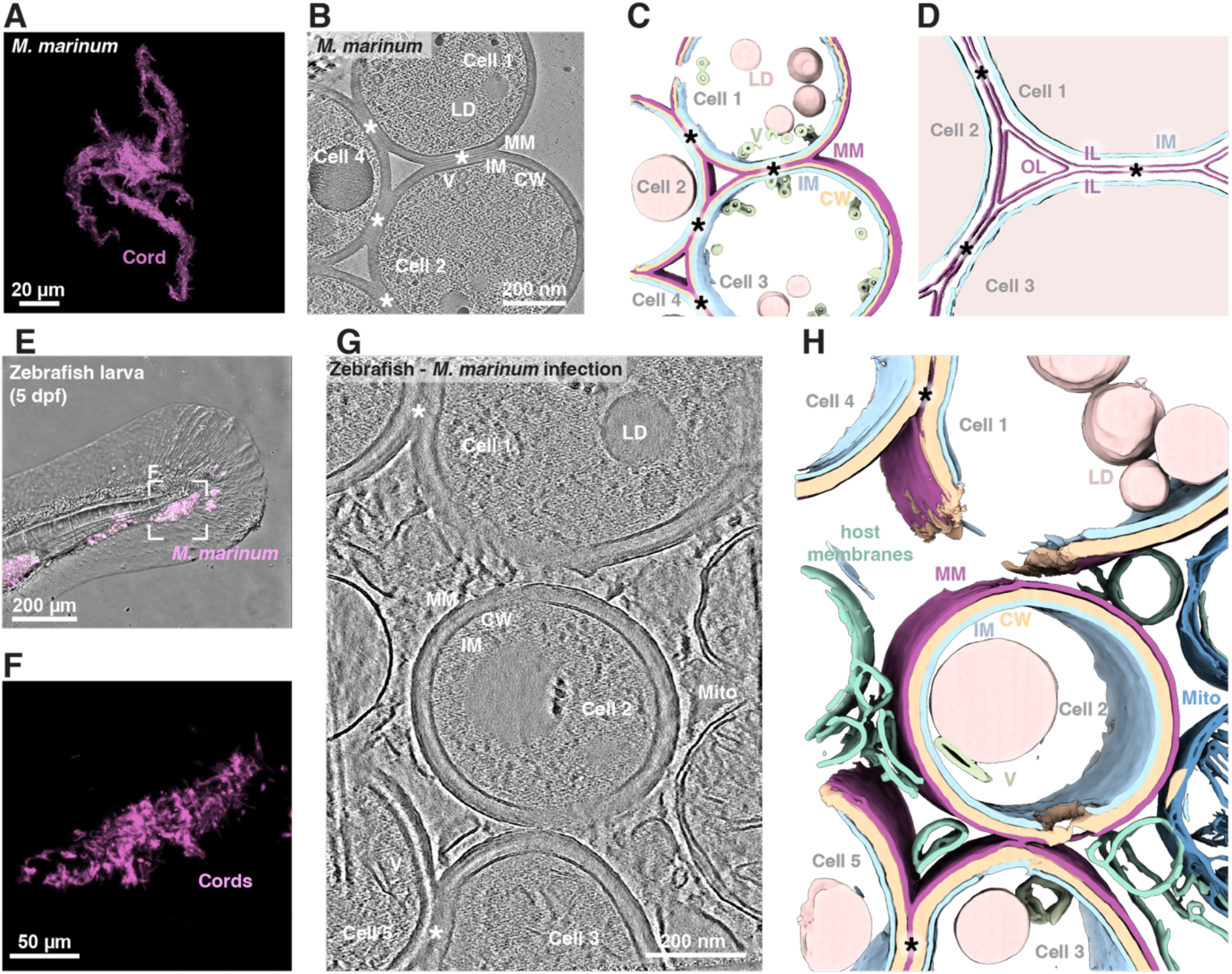
Mycobacterial cell-cell contacts in infected tissue share a continuous mycomembrane. (A) Fluorescence light microscopy of *M. marinum* cords. (B-D) Cryo-ET of *M. marinum* cord from axenic culture. (B) Tomographic slices through cord cell-cell contacts. (C-D) Three-dimensional segmentations of the tomogram shown in panel B. (E-F) *M. marinum* (pseudocolored in magenta) in a zebrafish larva at 5 days post fertilization and 3.5 days post infection, (E) overlaid on the brightfield image of an infected zebrafish fin and (F) fluorescence channel alone at a higher magnification, highlighting mycobacterial cording. (G) Tomographic slice and (H) three-dimensional segmentation of mycobacteria in infected tissue with host membranes (green) and mitochondria (dark blue) surround bacteria.

To determine if cords forming during infection have the same architecture as was observed during axenic culture, we performed cryo-ET of cording *M. marinum* in infected zebrafish larvae (Figure S1F; Fig 2E-F). Tomograms of infected tissue showed bacteria embedded directly in host tissue, with host-derived membranes and mitochondria adjacent to the bacterial surface (Figure 2G and H). As in axenic culture, neighboring bacteria in infected tissue were directly apposed with no gaps between the envelopes. The mycomembranes were continuous across the contact, forming the same shared membrane comprising the inner leaflets of each cell, with a thickness of 10.9 nm (Fig 2G and H).

Thus, taking advantage of the *M. marinum*-zebrafish TB model, we were able to confirm that mycobacterial cording in vivo is also associated with membrane enmeshment. The finding that this striking architecture is robust across very different growth environments (bacterial growth media versus cellular debris) suggests that it is not an artifact of axenic growth.

### Membrane enmeshment occurs in non-cording *Mycobacteriales*

To determine if membrane enmeshment was a feature specific to cording mycobacteria, we examined *M. smegmatis*, an environmental mycobacterium, which forms disordered clumps rather than cords (Figure 3A). Imaging by cryo-ET revealed that apposed *M. smegmatis* cells also exhibited mycomembrane enmeshment, characterized by exclusion of the outer mycomembrane leaflets and the formation of a new membrane interface comprising the two inner leaflets of the adjoining cells (Figure 3B-E; Fig S2C). Thus, the presence of enmeshed interfaces is a shared feature of cording and clumping in mycobacteria.

**Figure 3.**
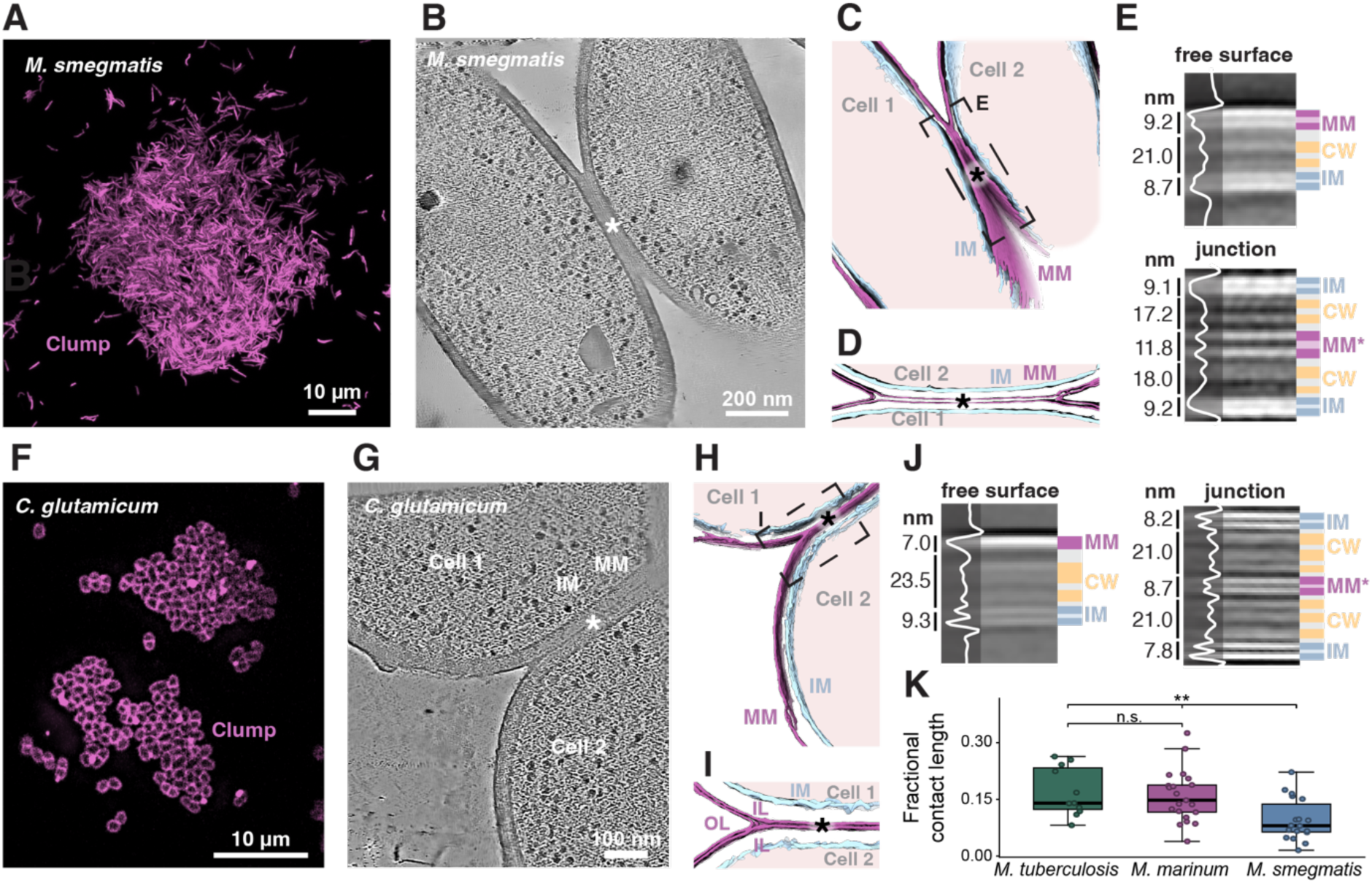
Mycomembrane enmeshment is not restricted to cording species. (A) Fluorescence microscopy of a *M. smegmatis* clump. (B-E) Cryo-ET of *M. smegmatis* in a clumped multicellular colony. (B) Tomographic slice through an *M. smegmatis* clump. (C-D) Segmentations of the tomogram shown in panel B. (E) Subtomogram analysis and density profiles across the free envelope (top) and the junction (bottom), with layers of the cell envelope marked. (F) Fluorescence light microscopy of a *C. glutamicum* clump (stained with Nile red). (G-J) Cryo-ET of *C. glutamicum* in a clumped multicellular colony. (G) Tomographic slice through a *C. glutamicum* clump. (H-I) Segmentations of *C. glutamicum* cell-cell contacts. (J) Subtomogram analysis and density profiles as in panel E. (K) Cell contact length of *M. tuberculosis* and *M. marinum* cords compared with *M. smegmatis* clumps normalized to cell perimeter. Box plots show median, with each point representing one tomogram. n=11 (*M. tuberculosis*), n=21 (*M. marinum*) and 18 (*M. smegmatis*). Comparisons are two-sided Mann-Whitney U tests on per-tomogram values. n.s.: not significant; \*\**p*<0.01.

We hypothesized that enmeshment might be linked to the mycolic-acid rich mycomembrane shared by mycobacteria. To probe this further, we tested *Corynebacterium glutamicum,* a non-pathogenic, non-mycobacterial member of *Mycobacteriales,* which possesses a mycomembrane and forms small, disorganized clumps (25). *C. glutamicum* revealed similar membrane enmeshment at areas of cellular contact (Figure 3F-J). These findings suggested that the capacity for enmeshment arises from shared envelope properties, likely those of mycomembranes.

While corded and clumped mycobacteria both showed enmeshed membranes, further analysis of cell-cell contacts showed that the length of the enmeshed regions in apposed *M. smegmatis* cells was less than those of *M. tuberculosis* and *M. marinum* (Fig 3L). These findings are consistent with a model in which enmeshment occurs as a consequence of mycomembrane properties shared by cording and noncording *Mycobacteriales.* Additional factors present in *M. tuberculosis* and *M. marinum* enable longer enmeshments.

### Mycomembrane enmeshments arise during cell division

Does enmeshment generate or stabilize cell association, or does stable cell association generate enmeshment? One possibility was that enmeshment forms at division, retaining daughters together and initiating clonal aggregation; cord-formers then extend and re-orient the inherited interfaces. Alternatively, cells might first become persistently apposed through another mechanism, with enmeshment subsequently forming at these contacts. The characteristic mode of division in *C. glutamicum* favored the first possibility. *C. glutamicum* divides by “V-snapping,” in which newly-divided daughters remain attached as characteristic V-shaped pairs, with repeated divisions generating angular clusters (26, 27). We asked whether both clumped and corded mycobacterial communities also arose through retention of daughter cell contacts generated at division. We used fluorescence microscopy to follow single *M. smegmatis, M. tuberculosis,* and *M. marinum* cells as they grew into communities (Figures 4A–C). In all three species, division left daughter cells joined at their poles in V- shaped intermediates, as described for *C. glutamicum* (27). In *M. smegmatis*, daughter cells continued growing in different directions, and repeated divisions generated disordered clumps of cells that remained attached (Figure 4A). In *M. tuberculosis* and *M. marinum*, daughter cells instead moved into alignment along their long axes, either by closure of the V-shaped pair or by sliding along one another to extend the contact (Figures 4B and C). Both routes produced aligned pairs that grew into corded microcolonies after repeated divisions. We quantified this divergence by following multiple divisions and measuring local alignment over time (Figure S3A-D). Local alignment in *M. smegmatis* communities remained limited as the clumps grew, whereas *M. marinum* colonies elongated their alignment and became increasingly enriched in highly ordered arrangements (Figure S3B and D). These findings showed that both clumps and cords could develop from single founder cells, with daughter cells remaining associated from division through subsequent community growth.

**Figure 4.**
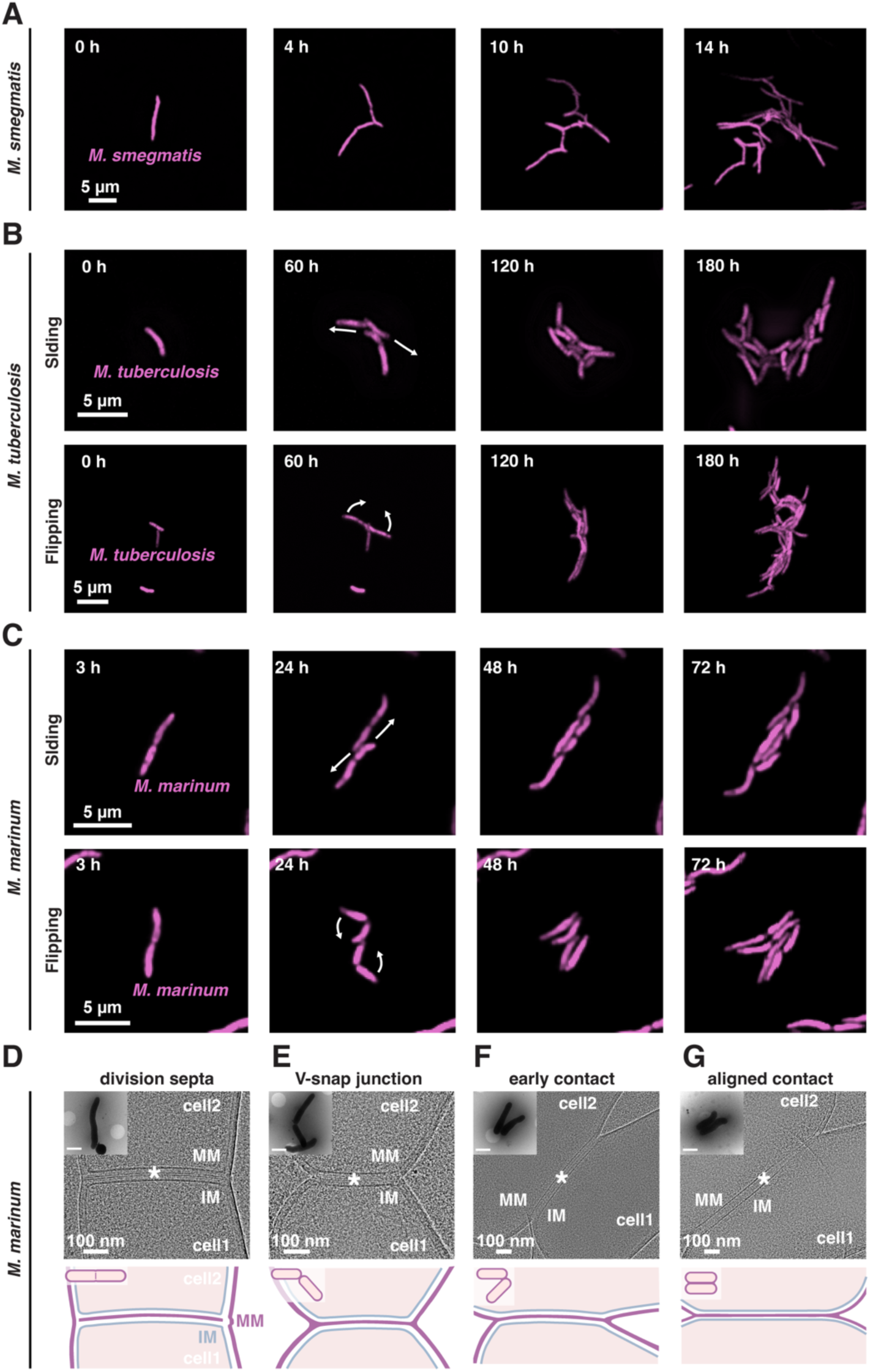
The enmeshed cell-cell junction forms at division and is retained as daughter cells grow into cords or non-aligned clumps. (A) Time-lapse fluorescence microscopy of *M. smegmatis* microcolony growth from a single cell over 14 h. (B) Time-lapse fluorescence microscopy of *M. tuberculosis* over 180 h, showing two modes of post-divisional cell-cell rearrangement: sliding, in which daughters move along the contact (top) and flipping (bottom), in which one daughter rotates about the division site. (C) As in B, live cell imaging of *M. marinum* (pseudocolored in magenta) over 72 h showing similar modes of post-divisional cell-cell rearrangement, sliding (top) and flipping (bottom). (D-G) Low-magnification cryo-EM of *M. marinum* at different stages of division: (D) formation of the division septum; (E) V-shaped cell pair post-division; (F) early and (G) late stage in the extension of the cell-cell contact. Enmeshed mycomembrane contacts highlighted with asterisks. Scale bars in insets, 1 µm.

To determine whether these division-generated contacts represented mycomembrane enmeshments, and whether the junctions persisted as daughter cells aligned, we imaged *M. marinum* cells at successive stages of division by cryo-ET (Figures 4D–G). We observed a mycomembrane lining the division septum, distinct from the mycomembrane lining the flanks of the emerging daughter cells (Figure 4D). In V-shaped daughter pairs, this septal mycomembrane was continuous with the surface mycomembrane at the new poles, forming an enmeshed mycomembrane junction (Figure 4E). Enmeshed junctions were also present between aligned daughter-cell pairs (Figures 4F and G). Across the division stages examined, we did not observe daughter cells separated by two distinct, free mycomembranes.

Together, these findings show that mycomembrane enmeshment originates during cell division, rather than arising only after previously separated cells come into contact. They support a model in which mycomembrane junctions present at cell division result in daughter cell retention and seeding of clonal communities. In *M. smegmatis*, these contacts remain in largely disordered arrangements, whereas in *M. tuberculosis and M. marinum* they are extended and reoriented as cells align into cords. Thus, enmeshment provides a common division-generated architecture that additional factors remodel in cord-forming mycobacteria.

### The mycomembrane lipid PDIM lengthens membrane enmeshments to enable cord formation

What factor(s) might enable interface extension and reorientation in cord-forming mycobacteria? Because membrane enmeshment appeared to be a shared mycomembrane property, we considered mycomembrane lipid candidates. One possible candidate was trehalose 6,6′-dimycolate (TDM), historically termed “cord factor,” which promotes cording in both *M. tuberculosis* and *M. marinum* (2) (Figures 5A and B; Figure S4). However, TDM seemed unlikely to be the distinguishing factor because it is also present in non-cording *M. smegmatis*, which possesses a functional homolog of PcaA, the enzyme responsible for proximal cis-cyclopropanation of α-mycolates. Indeed, this homolog can restore both mycolate cyclopropanation and cording in a *Mycobacterium bovis* BCG *pcaA* mutant (28). Nevertheless, given TDM’s longstanding association with cording, we examined whether PcaA-dependent cyclopropanation was required for extension of membrane enmeshments. Deletion of *pcaA* caused a relatively modest reduction in cording morphology in both *M. tuberculosis* and *M. marinum* (Figures 5A and B; Figure S4). Moreover, cryo-ET of the *M. marinum ΔpcaA* mutant revealed enmeshed cell-cell junctions of similar lengths to wild type (Figures 5C–F). Thus, although PcaA-dependent mycolate cyclopropanation contributes to cording morphology, it is not required for extension of enmeshments, suggesting that its effects on community organization occur through a distinct mechanism.

**Figure 5.**
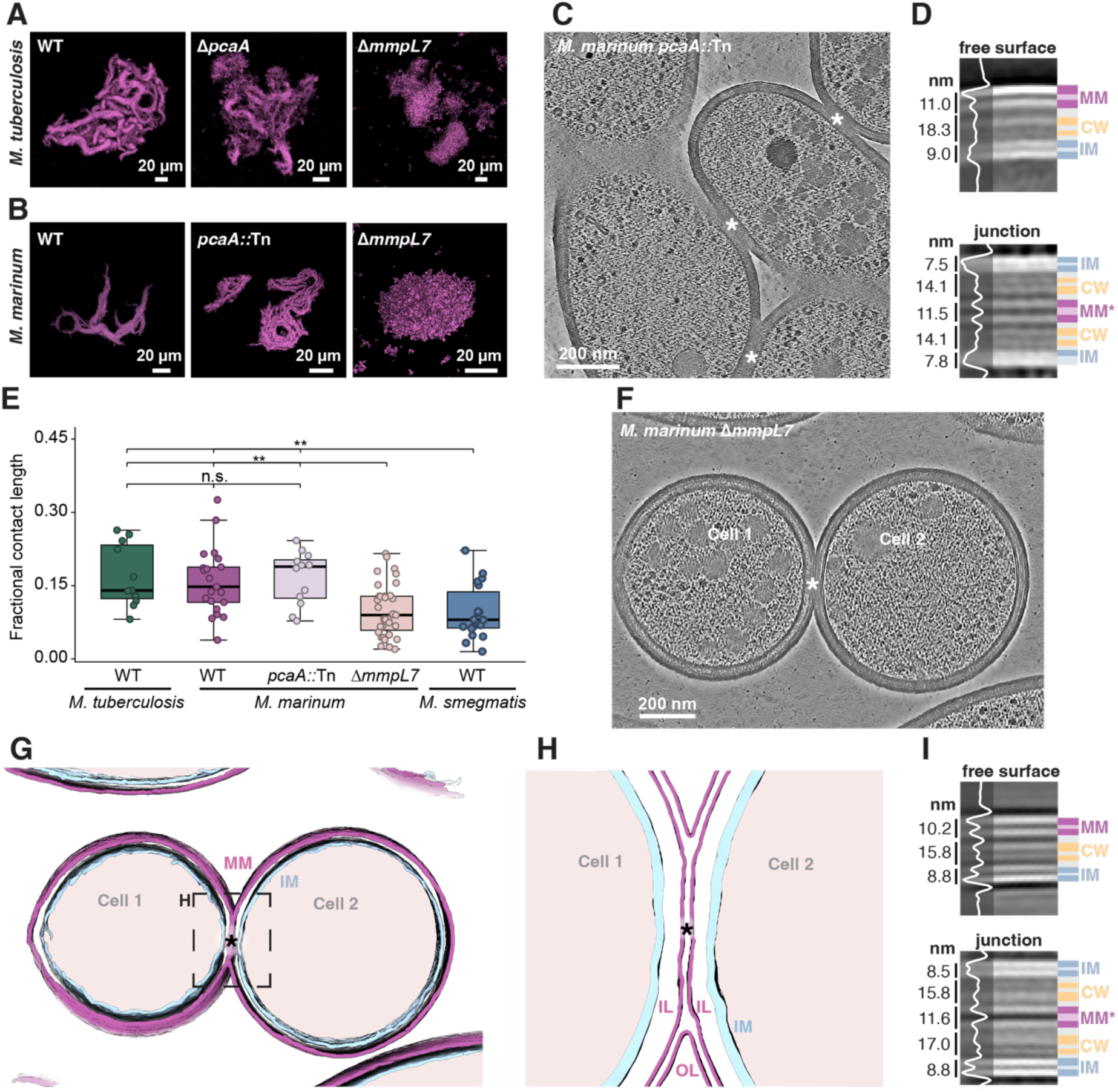
The virulence lipid PDIM extends mycomembrane enmeshment and is required for cord formation. (A-B) Fluorescence microscopy of (A) *M. tuberculosis* and (B) *M. marinum* wild type (WT; left), mutants of the *pcaA* gene (center; *M. tuberculosis*: Δ*pcaA*; *M. marinum*: *pcaA*::Tn) and strains lacking the *mmpL7* gene (right; *ΔmmpL7*). (C-D) Cryo-ET of *M. marinum pcaA*::Tn showing (C) tomographic slice and (D) subtomogram analysis across the free envelope and the junction. (E) Cell-cell contact length normalized to cell perimeter. Box plots show median, with each point representing one tomogram. n=11 (*M. tuberculosis*), n=21 (*M. marinum*), n=12 (*pcaA*::Tn), n=35 (Δ*mmpL7*) and n=18 (*M. smegmatis*). Comparisons are two-sided Mann-Whitney U tests on per-tomogram values. n.s.: not significant; \*\**p*<0.01. (F- I) Cryo-ET of *M. marinum* Δ*mmpL7*. (F) Tomographic slice, (G–H) envelope and junction segmentation and (I) subtomogram analysis across the free envelope and the junction.

The mycomembrane lipids PDIM seemed stronger candidates for enabling enmeshment extension in *M. tuberculosis* and *M. marinum*. PDIM biosynthetic capacity arose in the common ancestor of *M. marinum* and *M. tuberculosis* (29), and PDIM are present in both species but absent from the non-cording *M. smegmatis* and *C. glutamicum* examined here. Strikingly, loss of surface PDIM through mutation of its MmpL7 transporter caused a a loss of cording as expected (16), which we found to be more severe than in the corresponding *M. tuberculosis and M. marinum pcaA* mutants (Figures 5A and B; Figure S4). Cryo-ET of *M. marinum ΔmmpL7* showed that this failure to cord was associated with markedly reduced enmeshment lengths, comparable to those in *M. smegmatis* (Figures 5E–J; Figure S2D). In many cases, closely apposed cells lacked enmeshed interfaces altogether (Figure S2D).

Together, these findings indicate that surface PDIM is required for extension of mycomembrane enmeshments that enable the organization of clumps into cords.

### Membrane enmeshments create a shared mycomembrane between mycobacteria in cords

The topology of the cords revealed that the mycomembrane outer leaflet wrapped around the enmeshed bacteria and suggested that a shared mycomembrane compartment might have formed as a result. If so, we might expect that outer leaflet mycomembrane lipids traverse freely through the cords. To test this, we studied transfer of fluorescently labeled phenolic glycolipids (PGL) across the envelopes of corded *M. marinum,* using fluorescence recovery after photobleaching (FRAP) experiments. First, we photobleached the fluorophores on the entire surface of a single cell within a cord and observed nearly full recovery of the fluorescence intensity over time (t1/2=4.5 minutes (Figure 6A-C and Figure S5A-C)), suggesting that labeled PGL molecules were moving from adjoining areas of the cord into the bleached area. Control cells, that were not in a cord on the other hand, could not recover their fluorescence (Figure 6A-C). Next, we performed fluorescence loss in photobleaching (FLIP) experiments, repeatedly bleaching one area of a cord, which led to dimming of the rest of the cord (Figure 6D-E and Figure S5E-F). This confirmed that PGL could move between the corded cells. As a control, a standalone cell that was not bleached repeatedly suffered no comparable loss of fluorescence intensity in the same experiment (Figure 6D-E). These data confirmed that PGL could move between the corded cells.

**Figure 6.**
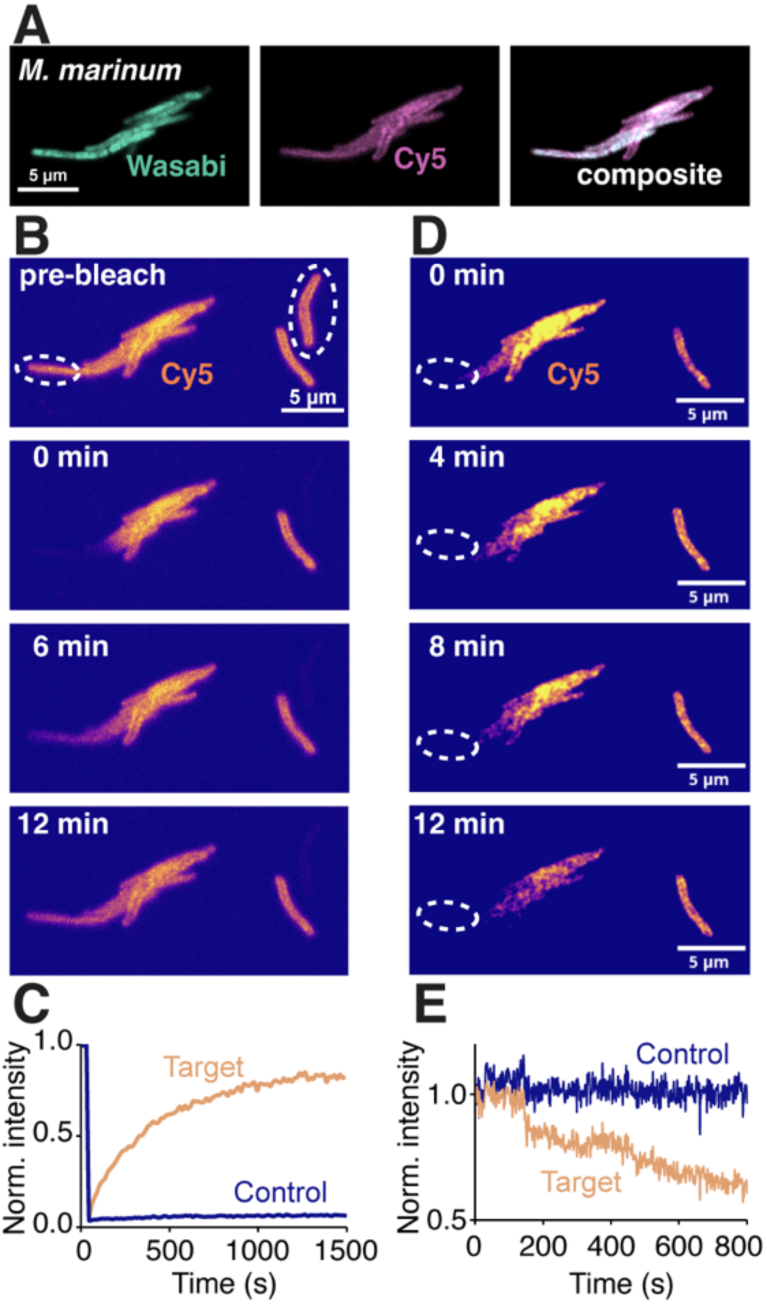
Phenolic glycolipid (PGL) diffuses through the shared mycomembrane compartment. (A) Early *M. marinum* (Wasabi, green, left) wild-type cords labelled with Cy5-PGL (magenta, center) and the composite (right). (B-C) Fluorescence recovery after photobleaching (FRAP). (B) A region within a cord (target) and a separate, non-contacting cell (control) were bleached and imaged over 12 minutes. Dashed outlines depict bleached regions. (C) Normalized fluorescence recovery for target and control regions. A single representative is shown; recovery was observed in all cords tested (n=13). (D-E) Fluorescence loss in photobleaching (FLIP). (D) Continuous bleaching of one region of a cord, with a non-contacting cell in the same field. (E) Normalized intensity during continuous bleaching. A single representative experiment is shown; fluorescence loss was observed in all cords tested (n=4).

Together, the FRAP and FLIP experiments showed that fluorescent PGL was not confined to individual cell envelopes but underwent long-range movement between cells within cords. Combined with our cryo-ET data that show a continuous outer leaflet of the mycomembrane between the cells in the cord, these data suggest that corded cells form a functionally connected lipid-exchange network.

## Discussion

Our findings suggest that mycobacterial community formation begins with a division-generated interbacterial mycomembrane interface. Non-cording species retain short enmeshments and form disordered clumps, whereas in *M. tuberculosis* and *M. marinum*, these interfaces extend and the cells become longitudinally aligned into cords. A striking feature of mycobacterial cords is the conspicuous absence of filamentous extracellular fibers of the kind observed in biofilm matrices formed by many other bacterial species (20). In these conventional biofilms, proteinaceous or polysaccharide fibers fulfil the dual roles of physically crosslinking cells (30) and embedding them within a protective extracellular scaffold (19). In contrast, in *M. tuberculosis* and *M. marinum*, cells are connected directly through mycomembrane enmeshment, without discernible intervening matrix material (Figure 7). Importantly, we found the same membrane architecture in *M. marinum* cords during zebrafish infection, establishing that these unusual interbacterial connections are relevant during infection, where the link between cording and virulence is relevant.

**Figure 7.**
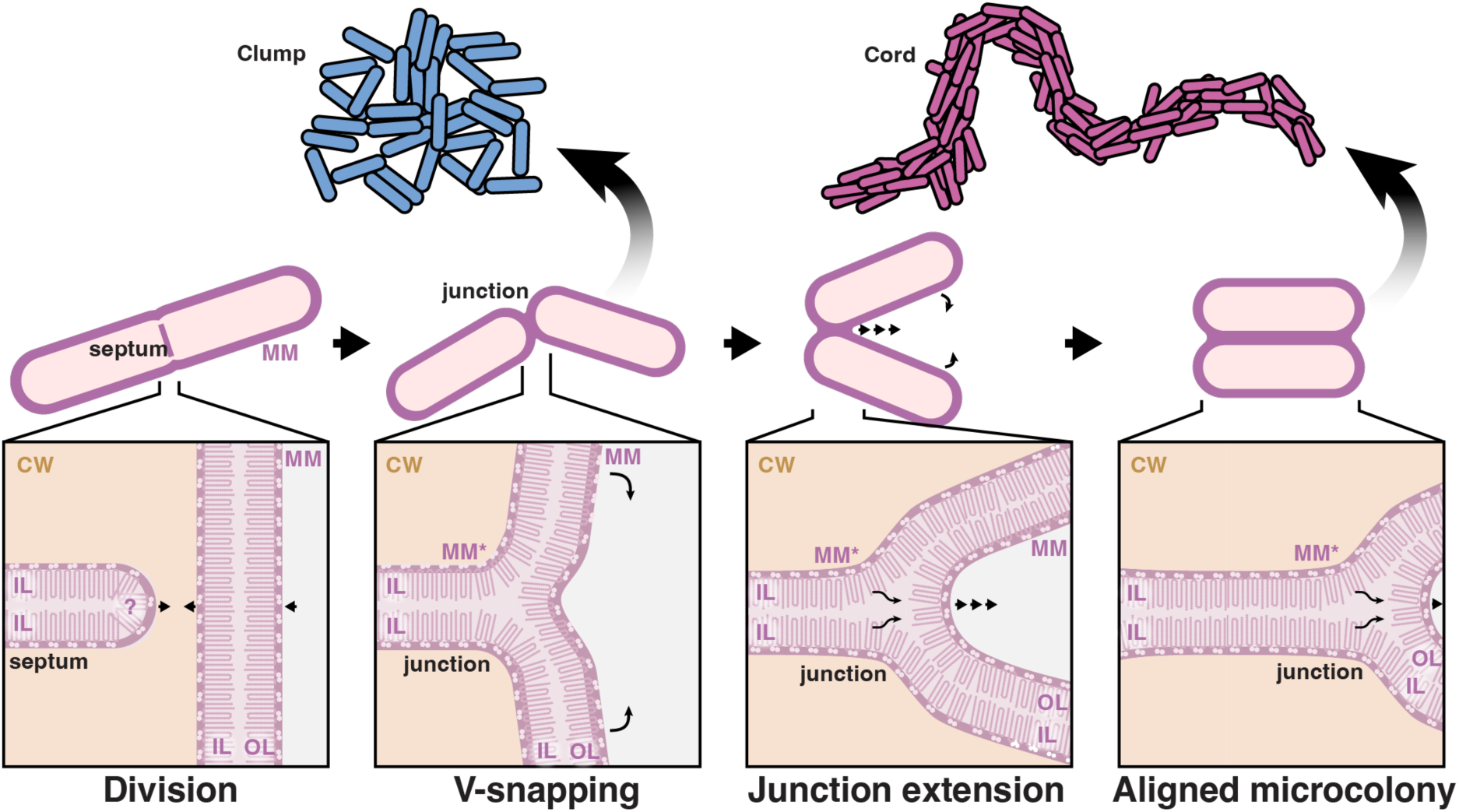
Proposed sequence from cell division to clump or cord formation. Top: community types; middle: cellular stages; bottom: magnified view of the cell envelope at the site of junction formation. *Division:* a septum containing a precursor mycomembrane forms between the two future daughter cells. *V-snapping:* daughters snap apart at the division site but remain physically linked by a shared mycomembrane (MM*), in which the inner leaflets of both cells form a single bilayer spanning the junction. *Junction extension and alignment:* cording species extend the junction along the cell sides (triple arrows), drawing sister cells into side-by-side contact (curved arrows) and, over successive divisions, building aligned microcolonies (right); in *M. marinum* and *M. tuberculosis*, this step requires phthiocerol dimycocerosate (PDIM). Clumping species do not extend the junction, so sister cells remain V-snapped and are arranged randomly by growth, forming disorganized clumps (left). CW: cell wall; MM: mycomembrane; MM*: shared mycomembrane at the junction; IL and OL: inner and outer leaflet of the mycomembrane.

From a mechanistic standpoint, the observation that enmeshed contact sites are established during cell division suggests a model in which a nascent patch of shared membrane is inherited at the division septum and subsequently expanded in cording species. Thus, extensive enmeshment can develop without requiring de novo fusion events between already separated cells. In *M. marinum*, we found that surface localization of the mycomembrane lipids PDIM is required for extension of these post-division enmeshments, which is needed for cording. PDIM are large, highly apolar, methyl-branched lipids of the mycomembrane outer leaflet that contribute to its permeability barrier (31, 32). Their unusual conical molecular geometry is thought to favor highly curved, non-bilayer membrane arrangements (33). This suggests a model in which PDIM promotes rearrangement of the outer leaflets at the boundaries of enmeshed interfaces, permitting their extension. Whether this is the case, and how it relates to the known biophysical properties of PDIM, remains to be determined.

A limitation of our study is that while we and others have found that PDIM surface localization is required for *M. tuberculosis* cording, we have not directly established that promotes extension of membrane enmeshments in this species. Notably, PDIM is not universally required for cording. *Mycobacterium abscessus*, which is more distantly related to *M. tuberculosis* and *M. marinum* and lacks PDIM, also forms cords both in axenic culture and in vivo during zebrafish infection (34). In *M. abscessus*, cord formation is linked to the *loss* of cell-surface lipids, the abundant glycopeptidolipids (GPLs), which is accompanied by a switch from smooth to rough colonial morphology (35, 36), mirroring the transition to smooth morphologies in *M. tuberculosis* strains with reduced PDIM expression (37). We do not yet know whether GPL loss in *M. abscessus* also promotes cord formation through extension of existing mycomembrane enmeshments. If so, it would be intriguing that two distinct changes in mycomembrane lipid composition -acquisition of PDIM in *M. tuberculosis* and *M. marinum* and loss of GPLs in *M. abscessus* - can promote the same membrane architecture. Alternatively, these species may form morphologically similar cords with distinct underlying architecture.

Another unusual property of PDIM is its ability to move from mycobacterial into host-cell membranes (18, 33). In this connection, our finding that fluorescent phenolic glycolipid (PGL), which shares the dimycocerosate lipid scaffold with PDIM, moves freely throughout cords is intriguing. Because PDIM can transfer between separate membranes (18, 33), this observation does not necessarily establish diffusion through a single continuous shared membrane compartment. Nevertheless, it shows that lipid pools of neighboring bacteria within cords are dynamically connected, raising the possibility that such exchange has additional functions in these multicellular communities.

Our work uncovers a new function of PDIM. In addition to contributing to the intrinsic cell-envelope permeability barrier and promoting virulence through modulation of mycobacterium-macrophage interactions (14–18). PDIM promotes the interbacterial membrane interactions that play a pivotal role in cording. Future work will be needed to determine how PDIM enables or stabilizes extension of enmeshed interfaces, how it interacts with the mycomembrane factors that initiate enmeshment, and also how TDM contributes to cording. More broadly, our findings reveal that the mycomembrane is not simply a barrier surrounding individual bacteria but also provides the structural basis for clonal communities, whose organization into cords is determined by changes in envelope lipid composition.

## Materials and Methods

### Bacterial strains and cultivation

*M. marinum* M strain (ATCC BAA-535) and its mutant derivatives Δ*mmpL7* and *pcaA*::Tn expressing BFP2, mWasabi or tdTomato under the control of the *msp12* promoter were used in this study, together with *M. tuberculosis* Δ*leuD* Δ*panCD* mc^2^ 6206 and its mutant derivatives *ΔpcaA* and *ΔmmpL7* expressing *msp12:tdTomato*, *M. smegmatis* mc^2^ 155 expressing *msp12:tdTomato* and *C. glutamicum* strain 541 (ATCC 13058). Mycobacteria were grown in Middlebrook 7H9 medium at 37 C (*M. tuberculosis, M. smegmatis*) or 33 °C (*M. marinum*) under antibiotic selection as appropriate. *C. glutamicum* was grown in beef-peptone medium at 37 °C. For media composition, supplements and selection and for expanded protocols on the generation of the *M. tuberculosis* Δ*mmpL7* strain, see SI Appendix, Supplementary Materials and Methods.

### Zebrafish husbandry and infections

Wild-type AB (Zebrafish International Resource Center, ZIRC) and *Tg(mpeg1:Brainbow)^w201^*(*mpeg1:tdTomato*) zebrafish were maintained under standard conditions. Embryos were raised at 28.5 °C and were transferred to 0.5× E2 medium supplemented with 0.003% PTU (1-phenyl-2-thiourea (Sigma)) at 1 day post fertilisation (dpf) to inhibit melanin synthesis. For infections, 1.5 dpf larvae were dechorionated with ≤0.5 mg/mL pronase (Sigma-Aldrich) and then anesthetized with fish water containing 0.025% tricaine (Sigma). Larvae were injected via the caudal vein or the fin using single-cell suspensions of *M. marinum* of known titer to deliver 100–250 bacteria per 3–5 nanolitre injection. Phenol red sodium salt (≤ 1% w/v diluted in PBS, Sigma-Aldrich) was used as an injection tracer. Inoculums were confirmed by injecting onto Middlebrook 7H10 agar plates (supplemented with OADS, plus hygromycin B or kanamycin, as appropriate). See SI Appendix, Supplementary Materials and Methods for husbandry, embryo medium composition and the generation of single cell suspensions.

### Cryogenic sample preparation and cryo-FIB milling of cultured bacteria

Bacteria were grown to a density of OD600 ∼1.0 in Middlebrook 7H9 medium without Tween, concentrated 20-fold and gently resuspended in Middlebrook 7H9 medium supplemented with 10% (v/v) glycerol. *M. smegmatis* and *M. marinum* clumps and cords were applied to glow-discharged Quantifoil R1/4 silicon dioxide 200-mesh gold grids and were vitrified by plunge-freezing into liquid ethane using a Thermo Scientific Vitrobot Mark IV with excess liquid removed by backside blotting. *M. tuberculosis* cords were applied to Quantifoil R1.2/1.3 holey carbon 200-mesh gold grid and vitrified by high-pressure freezing (HPF) on a Compact 03 HPF (M. Wohlwend) following the ‘waffle’ method (38). Prior to lamella cryo-FIB milling, *M. tuberculosis* cell clusters were localized by cryo-fluorescence microscopy. Lamellae were prepared on a Zeiss Crossbeam 550 field emission scanning electron microscope equipped with a Quorum cryo-stage, with the specimen chamber and stage maintained below –160 °C. Plunge-frozen specimens were thinned by a structured, four-step milling sequence. Specimens vitrified by HPF were thinned by rough trench milling followed by gradual sequential thinning. For more details on sample vitrification, cryo-CLEM and cryo-lamella milling, see SI Appendix, Supplementary Materials and Methods.

### Cryogenic sample preparation of infected zebrafish tissues

Zebrafish larvae infected with *M. marinum* were processed at 3.5 days post infection. Anaesthetized larvae were micro-sectioned in 0.5x E2 supplemented with 0.025% (w/v) Tricaine and 10% (v/v) glycerol, transferred to a 2 mm × 1 mm copper slot grid, assembled into an HPF sandwich and HPF vitrified by HPF. Infection sites were located by cryo-fluorescence microscopy on a Zeiss LSM 900 equipped with a Linkam cryo-stage. Lamellae were generated as described above with minor adaptations. For the planchette assembly and detailed protocols, see SI Appendix, Supplementary Materials and Methods.

### Cryo-ET data collection

Tilt-series were collected at 300 kV on either a Titan Krios G3 equipped with a K3 Gatan detector behind a Gatan Imaging Filter (20 eV slit width) or a Titan Krios G4 equipped with a Falcon 4i camera behind a Selectris-X Energy filter (10 eV slit width) using the SerialEM software and a dose symmetric tilting scheme. Dataset-specific acquisition parameters are given in SI Appendix, Supplementary Materials and Methods and Table S1.

### Cryo-ET processing, segmentation and analysis

Tilt-series were processed in RELION5.0 (39). Movies were motion-corrected using the MotionCor2 (40) implementation, splitting the data into odd and even frame subsets for downstream processing. Tilt-series alignment and contrast transfer function (CTF) estimation were performed using ARETOMO2 (41). Three-dimensional tomograms were reconstructed at a final pixel size of 10 Å/pixel. Tomograms were denoised with cryo-CARE (42). Membranes were segmented using AIS (43) and subvolumes along membranes were extracted and analysed by subtomogram analysis as described previously (23). Cell-cell contact sites were quantified via manual segmentation using IMOD (44). Figures displaying tomographic slices and segmentations were prepared from denoised tomograms (43, 45), rendered using IMOD (44), Fiji (46) and UCSF ChimeraX (47) with ArtiaX plugin (48). Tilt series of *C. glutamicum* were originally acquired for (25). For this study, tilt series were reprocessed de novo from the raw frames as described above. See SI Appendix, Supplementary Materials and Methods for expanded protocols on cryo-ET data segmentation and analysis.

### Confocal light microscopy of whole colonies

Cultures were grown in 7H9 with OADS at 33 °C (*M. marinum*) or 37 °C (*M. tuberculosis* and *M. smegmatis*) to an OD600 of ∼1.0 and 5 μL was spotted onto a 1.5% agarose pad prepared in 7H9 with OADS in a glass-bottom confocal dish. *C. glutamicum*, which carries no fluorescent reporter, was stained with 1 μM Nile red for 10 min and washed with PBSbefore spotting. Colonies were imaged on a Zeiss LSM 710 confocal microscope with a 63× Plan-Apochromat oil-immersion objective.

### Live confocal time-lapse imaging

Single-cell suspensions of wild-type and mutant *M. marinum*, *M. smegmatis* and *M. tuberculosis* were spotted onto a 1.5% agarose pad made with 7H9 supplemented with OADS and hygromycin. Time-lapse imaging was conducted on an inverted Zeiss LSM 710 laser scanning confocal microscope with images acquired every 15 min for *M. smegmatis*, every 60 min for *M. marinum* and every 90 min for *M. tuberculosis.* Stacks were drift -orrected using Fiji SIFT (49). Colony organization was quantified with a custom Python script built on NumPy, SciPy and scikit-image (50). For more details, see SI Appendix, Supplementary Materials and Methods.

### Fluorescence recovery after photobleaching (FRAP) and fluorescence loss in photobleaching (FLIP)

For metabolic labeling of phenolic glycolipid (PGL), wild-type and mutant *M. marinum* were cultured in the presence of 750 μM 3-azido-4-hydroxybenzoic acid for 24h at 33°C, washed and labeled with 25 μM membrane-impermeant sulfo-Cy5-DBCO for 30 min at room temperature. Cells were washed and spotted onto a 1.5% agarose pad made in a glass-bottom confocal dish. Photobleaching was performed on an inverted Zeiss LSM 710 laser scanning confocal microscope using the 633 nm laser line. For FRAP, an entire cell was photobleached and recovery was recorded at 15 s intervals over 15 min. For FLIP, one cell was repeatedly photobleached over 15 min. Image stacks were registered and background-subtracted mean fluorescence intensity of each region of interest was extracted for every frame using a custom Fiji macro. See SI Appendix Supplementary Materials and Methods for more details.

## Materials availability

All unique/stable reagents generated in this study are available without restriction from the corresponding authors.

## Data availability

The cryo-ET tomograms will be deposited at the Electron Microscopy Data Bank before publication.

## Acknowledgments

This work was supported by the Swiss National Science Foundation Postdoc Mobility fellowship P500PB_225494 and by European Molecular Biology Organization Postdoctoral fellowship ALTF 1283-2025 (K.J.), a Wellcome Trust Principal Research Fellowship (223103/Z/21/Z) and National Institutes of Health R01AI054503 (L.R) and the Medical Research Council, as part of United Kingdom Research and Innovation (also known as UK Research and Innovation) Program MC_UP_1201/31 the Wellcome Trust (Grant 225317/Z/22/Z), the Lister Institute for Preventative Medicine and the European Molecular Biology Organization (T.A.M.B.) We thank J. Shanahan for generating the *M. tuberculosis ΔmmpL7* mutant strain used in this study; B. Cormack, P. Edelstein, A. Fountain, J. Shanahan and O. Smith for discussion; B. Cormack, P. Edelstein and J. Shanahan for manuscript review; the LMB media kitchen; the LMB electron microscopy facility and the LMB computing facility, particularly T. Darling, J. Grimmett, and I. Clayson.

## Author Contributions

L.R. and T.A.M.B. conceived and supervised the project and acquired funding for it; K.J. performed experiments; K.J., L.R. and T.A.M.B. analysed and interpreted data and wrote the paper.

## References

1. R. Koch, Die Ätiologie der Tuberkulose (The Aetiology of Tuberculosis). Berliner klinische Wochenschrift 19, 221–230 (1882).

2. M. S. Glickman, J. S. Cox, W. R. Jacobs, A novel mycolic acid cyclopropane synthetase is required for cording, persistence, and virulence of Mycobacterium tuberculosis. Mol Cell 5, 717–727 (2000).

3. A. J. Pagán, L. Ramakrishnan, The Formation and Function of Granulomas. Annu. Rev. Immunol. 36, 639–665 (2018).

4. Y. S. McCarter, I. N. Ratkiewicz, A. Robinson, Cord formation in BACTEC medium is a reliable, rapid method for presumptive identification of Mycobacterium tuberculosis complex. J Clin Microbiol 36, 2769–2771 (1998).

5. A. J. Lenaerts, et al., Location of persisting mycobacteria in a Guinea pig model of tuberculosis revealed by r207910. Antimicrob Agents Chemother 51, 3338–3345 (2007).

6. L. Arias, et al., Cording Mycobacterium tuberculosis Bacilli Have a Key Role in the Progression towards Active Tuberculosis, Which is Stopped by Previous Immune Response. Microorganisms 8, 228 (2020).

7. R. Mishra, et al., Mechanopathology of biofilm-like Mycobacterium tuberculosis cords. Cell 186, 5135–5150.e28 (2023).

8. K. Abe, S. Ohtani, M. Hara, “Cord formation” in smear specimen prepared from sputum for a more rapid method of presumptive identification of Mycobacterium tuberculosis. Rinsho Byori 54, 116–120 (2006).

9. G. Middlebrook, R. J. Dubos, C. Pierce, Virulence and morphological characteristics of mammalian tubercle bacilli. J Exp Med 86, 175–184 (1947).

10. V. Jarlier, H. Nikaido, Mycobacterial cell wall: Structure and role in natural resistance to antibiotics. FEMS Microbiology Letters 123, 11–18 (1994).

11. P. J. Brennan, H. Nikaido, The envelope of mycobacteria. Annu Rev Biochem 64, 29–63 (1995).

12. H. Clay, H. E. Volkman, L. Ramakrishnan, Tumor necrosis factor signaling mediates resistance to mycobacteria by inhibiting bacterial growth and macrophage death. Immunity 29, 283–294 (2008).

13. D. M. Tobin, et al., The lta4h locus modulates susceptibility to mycobacterial infection in zebrafish and humans. Cell 140, 717–730 (2010).

14. J. S. Cox, B. Chen, M. McNeil, W. R. Jacobs, Complex lipid determines tissue-specific replication of Mycobacterium tuberculosis in mice. Nature 402, 79–83 (1999).

15. L. R. Camacho, D. Ensergueix, E. Perez, B. Gicquel, C. Guilhot, Identification of a virulence gene cluster of Mycobacterium tuberculosis by signature-tagged transposon mutagenesis. Mol Microbiol 34, 257–267 (1999).

16. J. Yu, et al., Both phthiocerol dimycocerosates and phenolic glycolipids are required for virulence of Mycobacterium marinum. Infect Immun 80, 1381–1389 (2012).

17. C. J. Cambier, et al., Mycobacteria manipulate macrophage recruitment through coordinated use of membrane lipids. Nature 505, 218–222 (2014).

18. C. J. Cambier, S. M. Banik, J. A. Buonomo, C. R. Bertozzi, Spreading of a mycobacterial cell-surface lipid into host epithelial membranes promotes infectivity. Elife 9, e60648 (2020).

19. J. Böhning, et al., Donor-strand exchange drives assembly of the TasA scaffold in Bacillus subtilis biofilms. Nat Commun 13, 7082 (2022).

20. J. Böhning, A. K. Tarafder, T. A. M. Bharat, The role of filamentous matrix molecules in shaping the architecture and emergent properties of bacterial biofilms. Biochem J 481, 245–263 (2024).

21. G. R. Squyres, D. K. Newman, Single-cell lysis patterns morphogenesis of eDNA in the matrix of Pseudomonas aeruginosa biofilms. Proc Natl Acad Sci U S A 122, e2514210122 (2025).

22. V. Berk, et al., Molecular architecture and assembly principles of Vibrio cholerae biofilms. Science 337, 236–239 (2012).

23. T. A. M. Bharat, P. C. Hoffmann, W. Kukulski, Correlative Microscopy of Vitreous Sections Provides Insights into BAR-Domain Organization In Situ. Structure 26, 879–886.e3 (2018).

24. A. J. Fountain, et al., Structural and functional analysis of the Mycobacterium tuberculosis MmpS5L5 efflux pump presages increased bedaquiline resistance. Proc Natl Acad Sci U S A 122, e2516660122 (2025).

25. B. Isbilir, A. Yeates, V. Alva, T. A. M. Bharat, Mapping the ultrastructural topology of the corynebacterial cell surface. PLoS Biol 23, e3003130 (2025).

26. A. Neumeyer, T. Hübschmann, S. Müller, J. Frunzke, Monitoring of population dynamics of Corynebacterium glutamicum by multiparameter flow cytometry. Microb Biotechnol 6, 157–167 (2013).

27. X. Zhou, et al., Sequential assembly of the septal cell envelope prior to V snapping in Corynebacterium glutamicum. Nat Chem Biol 15, 221–231 (2019).

28. L. Alibaud, et al., Temperature-dependent regulation of mycolic acid cyclopropanation in saprophytic mycobacteria: role of the Mycobacterium smegmatis 1351 gene (MSMEG_1351) in CIS-cyclopropanation of alpha-mycolates. J Biol Chem 285, 21698–21707 (2010).

29. A. T. Vincent, et al., The Mycobacterial Cell Envelope: A Relict From the Past or the Result of Recent Evolution? Front Microbiol 9, 2341 (2018).

30. J. Böhning, et al., Architecture of the biofilm-associated archaic Chaperone-Usher pilus CupE from Pseudomonas aeruginosa. PLoS Pathog 19, e1011177 (2023).

31. L. R. Camacho, et al., Analysis of the Phthiocerol Dimycocerosate Locus ofMycobacterium tuberculosis. Journal of Biological Chemistry 276, 19845–19854 (2001).

32. C. V. Mulholland, et al., Propionate prevents loss of the PDIM virulence lipid in Mycobacterium tuberculosis. Nat Microbiol 9, 1607–1618 (2024).

33. J. Augenstreich, et al., The conical shape of DIM lipids promotes Mycobacterium tuberculosis infection of macrophages. Proc Natl Acad Sci U S A 116, 25649–25658 (2019).

34. I. Halloum, et al., Deletion of a dehydratase important for intracellular growth and cording renders rough Mycobacterium abscessus avirulent. Proc Natl Acad Sci U S A 113, E4228–4237 (2016).

35. S. T. Howard, et al., Spontaneous reversion of Mycobacterium abscessus from a smooth to a rough morphotype is associated with reduced expression of glycopeptidolipid and reacquisition of an invasive phenotype. Microbiology (Reading) 152, 1581–1590 (2006).

36. A. Bernut, et al., Mycobacterium abscessus cording prevents phagocytosis and promotes abscess formation. Proc Natl Acad Sci U S A 111, E943–952 (2014).

37. D. Giovannini, et al., A new Mycobacterium tuberculosis smooth colony reduces growth inside human macrophages and represses PDIM Operon gene expression. Does an heterogeneous population exist in intracellular mycobacteria? Microb Pathog 53, 135–146 (2012).

38. K. Kelley, et al., Waffle Method: A general and flexible approach for improving throughput in FIB-milling. Nat Commun 13, 1857 (2022).

39. A. Burt, et al., An image processing pipeline for electron cryo-tomography in RELION- 5. FEBS Open Bio 14, 1788–1804 (2024).

40. S. Q. Zheng, et al., MotionCor2: anisotropic correction of beam-induced motion for improved cryo-electron microscopy. Nat Methods 14, 331–332 (2017).

41. S. Zheng, et al., AreTomo: An integrated software package for automated marker-free, motion-corrected cryo-electron tomographic alignment and reconstruction. J Struct Biol X 6, 100068 (2022).

42. T.-O. Buchholz, et al., Content-aware image restoration for electron microscopy. Methods Cell Biol 152, 277–289 (2019).

43. M. G. F. Last, L. Abendstein, L. M. Voortman, T. H. Sharp, Streamlining segmentation of cryo-electron tomography datasets with Ais. Elife 13, RP98552 (2024).

44. J. R. Kremer, D. N. Mastronarde, J. R. McIntosh, Computer visualization of three-dimensional image data using IMOD. J Struct Biol 116, 71–76 (1996).

45. M. G. F. So-Last, A. Burt, T. Hale, M. Allegretti, Easymode: general pretrained networks for cellular cryo-ET enable flexible approaches to subtomogram averaging. [Preprint] (2026). Available at: http://biorxiv.org/lookup/doi/10.64898/2026.05.19.726344 [Accessed 18 October 2026].

46. J. Schindelin, et al., Fiji: an open-source platform for biological-image analysis. Nat Methods 9, 676–682 (2012).

47. E. F. Pettersen, et al., UCSF ChimeraX: Structure visualization for researchers, educators, and developers. Protein Sci 30, 70–82 (2021).

48. U. H. Ermel, S. M. Arghittu, A. S. Frangakis, ArtiaX: An electron tomography toolbox for the interactive handling of sub-tomograms in UCSF ChimeraX. Protein Sci 31, e4472 (2022).

49. C. Paganelli, et al., Scale Invariant Feature Transform as feature tracking method in 4D imaging: a feasibility study. Annu Int Conf IEEE Eng Med Biol Soc 2012, 6543–6546 (2012).

50. S. van der Walt, et al., scikit-image: image processing in Python. PeerJ 2, e453 (2014).

