## Supporting Information for "Acquisition of phthiocerol dimycocerosates enabled extension of fused mycomembranes for cellular cording in *Mycobacterium tuberculosis*"

This PDF file includes:

SI Materials and Methods

Table S1

Figures S1 to S5

SI References

**Supplementary Materials and Methods**

**Bacterial strains and cultivation.** *M. marinum* M strain (ATCC BAA-535) and its mutant derivatives Δ*mmpL7* and *pcaA*::Tn expressing BFP2, mWasabi or tdTomato under the control of the *msp12* promoter were grown at 33 °C under 50 µg/mL hygromycin B (Cambridge Bioscience) or 25 µg/mL kanamycin (Sigma-Aldrich) selection in Middlebrook 7H9 medium (BD Difco) supplemented with sodium oleate, albumin, dextrose and NaCl (OADS). *M. tuberculosis* Δ*leuD* Δ*panCD* mc$^2$ 6206 and its mutant derivatives Δ*pcaA* and Δ*mmpL7* expressing *msp12:tdTomato* were grown at 37 °C under hygromycin B and kanamycin selection in Middlebrook 7H9 medium supplemented with sodium oleleate, albumin, dextrose, 0.05 mg/mL L-leucine and 0.024 mg/mL calcium pantothenate (Sigma-Aldrich). *M. smegmatis* mc$^2$ 155 expressing *msp12:tdTomato* was grown in 7H9 Middlebrook medium at 37 ˚C. *Corynebacterium glutamicum* strain 541 (ATCC 13058) was grown in beef-peptone medium at 37 ˚C. For expanded protocols on the generation of the *M. tuberculosis* Δ*mmpL7* strain, see SI Appendix, Supplementary Materials and Methods.

**Construction of *M. tuberculosis* Δ*mmpL7* mutant.** The *M. tuberculosis* Δ*mmpL7* strain was constructed by recombineering as previously described (1, 2). The recombineering plasmid was constructed by amplifying the upstream flanking region of mmpL7 (F primer: AACGACGGCCAGTGAATTACTTAAGGATATCGAATGTCCGGACGGAACC and R primer: GATCCACTAGTTCTAGAGCGGCCGCTCTGTGTAGACGGCCAGCCGGA) and the downstream flanking region of mmpL7 (F primer: ACATTATACGAAGTTATGGCGCGCCGCGTGGCTTCCGACCACG and R primer: CCATGATTACGCCAAGCTTGCATGCGATATCTTCCACAGTCAACATTCCCTT). Correct insertion of the deletion cassette was confirmed by sequencing PCR products of the region (F primer: TGACCTCGGCAACATACTGAC and R primer: CGGAAACCAGAAGTCGCACTG). All other steps were performed as described (2).

**Generation of single cell suspensions.** *M. marinum*, *M. tuberculosis* and *M. smegmatis* were grown to an optical density at 600 nm wavelength of light ($OD_{600}$) of 1.0 and pelleted at 3,000 × g for 10 min. 5 ml of bacterial culture in Middlebrook 7H9 medium were passed gently through a fine-gauge needle to disrupt clumps, followed by a low-speed spin at 100 × g for 10 min to pellet residual clumps. For live imaging, cells in the supernatant were pelleted at 3000 × g for 10 min, aliquoted and stored at –70°C. For infections of zebrafish larvae, the supernatant was gravity filtered through a 5 µm (microns) non-pyrogenic polyethersulfone membrane, single cells were pelleted at 3000 × g for 10 min, aliquoted and stored at –70 ˚C. The titer of bacterial single cell suspension was estimated by light microscopy and colony forming unit counts.

**Zebrafish husbandry and infections.** Zebrafish embryos and larvae of the wild-type AB strain (Zebrafish International Resource Center, ZIRC) *Tg(mpeg1:Brainbow)*$^{w201}$ (*mpeg1:tdTomato*) fluorescent reporter line was used in experiments. Zebrafish were maintained in buffered reverse osmotic water systems under a 14-hour light/10-hour dark cycle. Zebrafish larvae were fed paramecia twice daily, while juvenile and adult zebrafish were fed at least twice a day with dry food and brine shrimp. Zebrafish embryos were collected and cultured in reverse osmosis water containing 0.18 g/L Instant Ocean Salt supplemented with 0.25 µg/mL methylene blue at 28.5 °C. On 1 day post fertilisation (dpf), embryos to be used in experiments were transferred to 0.5× E2 medium (7.5 mM NaCl, 0.25 mM KCl, 0.5 mM $MgSO_4$, 0.075 mM $KH_2PO_4$, 0.025 mM $Na_2HPO_4$, 0.5 mM $CaCl_2$, and 0.35 mM $NaHCO_3$) supplemented with 0.003% PTU (1-phenyl-2-thiourea (Sigma)) to inhibit melanin synthesis.

For infections, 1.5 dpf larvae were dechorionated with ≤0.5 mg/mL pronase (Sigma-Aldrich) and then anesthetized with fish water containing 0.025% tricaine (Sigma). Larvae were injected via the caudal vein or the fin using single-cell suspensions of *M. marinum* of known titer to deliver 100–250 bacteria per 3–5 nanolitre injection. Phenol red sodium salt (≤ 1% w/v diluted in PBS, Sigma-Aldrich) was used as an injection tracer. Inoculums were confirmed by injecting onto Middlebrook 7H10 agar plates (supplemented with OADS, plus hygromycin B or kanamycin, as appropriate).

**Grid preparation by plunge freezing.** *M. smegmatis* and *M. marinum* clumps and cords were were grown to a density of $OD_{600}$ ~1.0 in Middlebrook 7H9 medium without Tween, concentrated 20-fold by pelleting at 3000 × g for 10 min and gently resuspended in Middlebrook 7H9 medium supplemented with 10% (v/v) glycerol and applied to glow-discharged Quantifoil R1/4 silicon dioxide 200-mesh gold grids and were vitrified by plunge-freezing into liquid ethane using a Thermo Scientific Vitrobot Mark IV maintained at 15 ˚C and 100% relative humidity. Excess liquid was removed by backside blotting prior to automated plunging.

**Grid preparation of *M. tuberculosis* by high-pressure freezing (HPF).** *M. tuberculosis* cords were vitrified by high-pressure freezing (HPF) using a Compact 03 HPF (M. Wohlwend) following the 'waffle' method protocol (3). In brief, a 3 µL aliquot of the bacterial sample was applied to a Quantifoil R1.2/1.3 holey carbon 200-mesh gold grid. The grid was sandwiched between the flat sides of two metal planchettes (3 mm diameter, 0.5 mm thickness). This assembly was loaded into an HPF holder (0.95 mm depth) and immediately subjected to HPF.

**Cryogenic fluorescence light microscopy (cryo-CLEM).** *M. tuberculosis* cell clusters within the high-pressure frozen specimen were localized by cryo-fluorescence microscopy using a Zeiss LSM 900 confocal microscope equipped with a Linkam cryo-stage. The cryo-stage was maintained at constant chamber and bridge temperatures of –190 ˚C.

**Cryo-Focused ion beam (cryo-FIB) milling.** Cryo-focused ion beam (cryo-FIB) milling was performed on a Zeiss Crossbeam 550 field emission scanning electron microscope equipped with a Quorum cryo-stage, with system control and patterning by SmartFIB software (v.1.15). The specimen chamber and stage were maintained at stable temperatures below –160 ˚C. Grids were first sputter-coated with platinum for 45 sat 5 mA, followed by deposition of organometallic platinum via the integrated gas injection system (GIS) for 60 s.

Cell clusters on plunge frozen grids were brought to the coincident point and milled at a nominal shallow tilt angle of 10° relative to the grid plane. Ion beam milling was executed at a fixed FIB accelerating voltage of 30 kV using a structured, four-step progressive step-down milling sequence using currents of 700 pA, 300 pA, 100 pA and 50 pA.

Lamellae of the high-pressure frozen *M. tuberculosis* cell clusters were generated sequentially by initial milling of rough trenches measuring 30 μm × 40 μm on both sides of the targeted regions perpendicular to the grid plane using a 30 nA FIB current. The targeted regions were gradually thinned down to a thickness of 5 μm using a step-down sequence of 7 nA, 3 nA and 1.5 nA FIB currents at a milling angle of 20°. Following the milling of a stress-release notch, final lamellae were produced using a four-step polishing sequence with currents of 700 pA, 300 pA, 100 pA and 50 pA.

**Cryogenic sample preparation of infected zebrafish tissues.** Zebrafish larvae infected with *M. marinum* were inoculated for 3.5 days in 0.5× E2 embryo medium. Prior to sectioning, infected larvae were anesthetized in E2/2 containing 0.025% (w/v) Tricane. Anaesthetized larvae were transferred to E2/2 supplemented with 10% (v/v) glycerol. Sorting and fluorescence-based localization of infection sites were performed under a Nikon SMZ18 with an integrated CoolLED pE-300 fluorescence illumination unit. Infection sites containing fluorescent bacteria within the larvae were micro-sectioned using a scalpel. HPF sandwich assemblies were prepared under the stereomicroscope. The micro-sectioned tissues were carefully transferred into a 2 mm × 1 mm copper slot grid positioned on top of a 3 mm diameter, 0.5 mm thick aluminum planchette pre-coated with 0.1% (w/v) soy lecithin in chloroform. A 3 mm diameter, 0.1 mm thick spacer ring was placed over the grid and the assembly was closed using a 3 mm diameter, 0.26 mm thick top planchette, also pre-coated with 0.1% soy lecithin. Finally, a 3 mm diameter, 0.1 mm thick top spacer ring was added to complete the carrier assembly. The fully assembled tissue sandwich was immediately vitrified by HPF using a Wohlwend Compact 03 system.

*M. marinum* cell clusters within infected tissue inside the high-pressure frozen specimen were localized by cryo-fluorescence microscopy using a Zeiss LSM 900 confocal microscope equipped with a Linkam cryo-stage. The cryo-stage was maintained at constant chamber and bridge temperatures of -190 ˚C.

Lamellae were then generated with some adaptations to the way described for *M. tuberculosis* suspensions above. In brief, grids were first sputter-coated repeatedly (4×) with platinum for 30 s at 5 mA with 15-min pauses between depositions, followed by five rounds of deposition of organometallic platinum via the integrated gas injection system (GIS) for 60 s with 30 s pauses between depositions. Milling was performed sequentially by initial milling of rough trenches measuring 30 µm × 40 µm on both sides of the targeted regions perpendicular to the grid plane using a 30 nA FIB current. The targeted regions were gradually thinned down to a thickness of 5 µm using a step sequence of 7 nA, 3 nA and 1.5 nA FIB currents at a milling angle of 20°. Following the milling of a stress-release notch, final lamellae were produced using a four-step polishing sequence with currents of 700 pA, 300 pA, 100 pA and 50 pA.

**Cryo-ET data collection.** Cryo-ET data was collected on either a Titan Krios G3 equipped with a K3 Gatan detector behind a Gatan Imaging Filter (20 eV slit width) or a Titan Krios G4 equipped with a Falcon 4i camera behind a Selectris-X Energy filter (10 eV slit width) using the SerialEM software and a dose symmetric tilting scheme at 300 kV. For the *M. tuberculosis* wild-type samples, datasets were acquired at a pixel size of 1.965 Å/pixel with a target defocus range of –6 to –10 µm, an oscillation range of ±60° in 3° increments and a total dose of 150 $e^-$/Å$^2$. For *M. marinum*, tilt-series were recorded at a calibrated pixel size of 3.015 Å/pixel with a target defocus range of –5 to –7 µm and an angular range of ±60° in 2° increments, yielding a cumulative electron dose of 90 $e^-$/Å$^2$. *M. smegmatis* wild-type, *M. marinum pcaA*::Tn and Δ*mmpL7* mutant strains were imaged using identical parameters but with an increased total dose of 120–150 $e^-$/Å$^2$ and defocus ranges of –4 to –6 µm and –5 to –7 µm. Data of zebrafish tissue infected with *M. marinum* was recorded at a pixel size of 2.128 Å/pixel, spanning a target defocus of –3 to –5 μm, an angular range of ±50° in 2° increments and a total cumulative dose of 100 $e^-/Å^2$.

**Cryo-ET data segmentation and analysis.** Automated semantic segmentation of membranes was performed using the AIS (4) framework. Initially, a small training set consisting of 50–100 manually curated tomographic training patches were used to train a baseline VGG-M convolutional neural network. This initial model was scaled for further training using the built-in ezm-2d network architecture.

To analyse membrane features by two-dimensional (2D) subtomogram analysis, the resulting binarized membrane segmentations were converted into discrete 3D coordinates and were imported into RELION5.0. Full subtomograms were extracted from CTF-corrected tomograms and the central three slices of each extracted subtomogram were Z-projected using a custom Python script and subjected to reference-free 2D classification within RELION5.0.

Cell-cell contact sites were quantified via manual segmentation using IMOD. To characterize cellular interactions, contacts were classified into two structural categories: 1) close contacts, which comprise all cell-cell interactions displaying a total envelope to envelope distance of less than 100 nm, encompassing both interfaces with visible intercellular gaps and those without; 2) enmeshed contacts, which comprise cell-cell contacts characterized by a complete absence of visible gap density between the interacting membrane envelopes. In addition, the total measured perimeter of all cells within the tomogram was recorded.

For presentation in figures, tomograms were denoised using self-supervised workflows for denoising with IsoNet (5) implemented in easymode (6). Figure panels displaying tomographic slices and 3D segmentation renderings were prepared using IMOD (7), Fiji (8) and UCSF ChimeraX (9) with ArtiaX plugin (10). Tilt series of *C. glutamicum* were originally acquired for (11). For this study, tilt series were reprocessed de novo from the raw frames as described above.

**Live confocal time-lapse imaging.** For live confocal time lapse imaging, wild-type and mutant *M. marinum*, *M. smegmatis* and *M. tuberculosis* single-cell suspension were spotted onto a 1.5% agarose pad made with 7H9 media supplemented with OADS and hygromycin. Time-lapse confocal imaging was conducted on an inverted Zeiss LSM 710 laser scanning confocal microscope (Carl Zeiss Inc.). mWasabi and tdTomato were excited using the 488 nm and 561 nm laser lines, respectively. Fluorescent emission was collected using spectral detectors set to

490–550 nm and 570–640 nm, with the pinhole set to 1.0 Airy Unit. The system was equipped with an environmental chamber maintained at 33°C for *M. marinum* or 37°C for *M. smegmatis* and *M. tuberculosis*. Images were acquired every 15 min for *M. smegmatis*, every 60 min for *M. marinum* or every 90 min for *M. tuberculosis* using ZEN software (Carl Zeiss Inc.). Time-lapse image stacks were drift corrected using Fiji SIFT (12). Colony organization was quantified with a custom Python pipeline built on NumPy, SciPy and scikit-image (13). Images were background-subtracted, median filtered and normalized. Colonies were segmented by Otsu thresholding, followed by mask smoothing and morphological closing. Small and boundary objects were removed before quantification. Local orientation within colonies was obtained from the 2D structure tensor of the fluorescence image, which returns the local direction of intensity elongation at every pixel. Alignment between neighboring cells was summarized as the nematic order parameter S2, calculated over 14 μm square windows tiled across each colony. S2 ranges from 0 for randomly oriented cells to 1 for perfectly aligned parallel cells. Each colony was assigned the median S2 of its window. The number of cells per colony was estimated by the median mask area of single cells.

**Fluorescence recovery after photobleaching (FRAP) and fluorescence loss in photobleaching (FLIP).** For metabolic labeling of phenolic glycolipid (PGL), wild-type and mutant *M. marinum* were cultured in 7H9 medium supplemented with OADS in the presence of 750 μM 3-azido-4-hydroxybenzoic acid for 24h at 33°C. Bacteria were washed three times with PBS and labeled with 25 μM membrane-impermeant sulfo-Cy5-DBCO for 30 min at room temperature. Unincorporated label was removed by washing three times with PBS before the cells were spotted onto a 1.5% agarose pad made with 7H9 medium and imaged in a glass-bottom confocal dish.

FRAP and FLIP were performed on an inverted Zeiss LSM 710 laser scanning confocal microscope, as described in previous sections. Cy5 was excited using the 633 nm laser line and fluorescent emission was collected at 650-720 nm with the pinhole set to 1.0 Airy Unit. For FRAP, after 3 pre-bleach scans, a single cell was photobleached using 100% bleaching power and 75 μs pixel dwell time and recovery was recorded at 15 s intervals over 15 min after photobleaching. For FLIP, one cell was subjected to repeated photobleaching over 15 min, while the fluorescence of the adjacent cells in the same field of view was monitored. In both experiments, a non-contacting cell was bleached in parallel as a non-recovery control and an unbleached reference cell and a cell-free background region were recorded for correction.

Image stacks were registered using Fiji SIFT and mean fluorescence intensity of each region of interest (ROI) was extracted for every frame using a custom Fiji macro. For FRAP, background subtracted intensities of bleached ROI were normalized to an unbleached reference cell and to the pre-bleach baseline. For FLIP, the normalized intensity of the connected neighbor was corrected against the non-contacting control cell and diffusive connectivity was expressed as the fractional loss of signal relative to baseline.

Table S1. Cryo-ET data collection.

| | *M. tuberculosis* WT | *M. marinum* WT | *M. marinum pcaA::*Tn | *M. marinum ΔmmpL7* | *M. smegmatis* WT | Zebrafish infected with *M. marinum* |
|---|---|---|---|---|---|---|
| Microscope | Titan Krios G4 | Titan Krios G4 | Titan Krios G4 | Titan Krios G4 | Titan Krios G4 | Titan Krios G3i |
| Detector | Falcon 4i | Falcon 4i | Falcon 4i | Falcon 4i | Falcon 4i | Gatan K3 |
| Software | SerialEM | SerialEM | SerialEM | SerialEM | SerialEM | SerialEM |
| Magnification | 64,000 x | 42,000 x | 42,000 x | 42,000 x | 42,000 x | 42,000 x |
| Voltage (kV) | 300 | 300 | 300 | 300 | 300 | 300 |
| Energy filter | Selectris-X | Selectris-X | Selectris-X | Selectris-X | Selectris-X | Gatan Imaging Filter |
| Slit width (eV) | 10 | 10 | 10 | 10 | 10 | 10 |
| Defocus range ($\mu$m) | -6 to -10 | -5 to -7 | -4 to -6 | -4 to -6 | -5 to -7 | -3 to -5 |
| Pixel size (Å) | 1.965 | 3.015 | 3.015 | 3.015 | 3.015 | 2.128 |
| Total exposure ($e^-$/Å$^2$) | 150 | 90 | 120 | 140 | 150 | 100 |
| Exposure per tilt ($e^-$/Å$^2$) | 2.5 | 1.5 | 1.9 | 2.3 | 2.5 | 2.0 |
| Tilt increments | 3° | 2° | 2° | 2° | 2° | 2° |
| Tilt-series scheme | dose-symmetrical | dose-symmetrical | dose-symmetrical | dose-symmetrical | dose-symmetrical | dose-symmetrical |
| Tilt range | ±60° | ±60° | ±60° | ±60° | ±60° | ±50° |
| Tilt series alignment | Aretomo2 | Aretomo2 | Aretomo2 | Aretomo2 | Aretomo2 | Aretomo2 |
| Tomogram reconstruction | RELION5.0 | RELION5.0 | RELION5.0 | RELION5.0 | RELION5.0 | RELION5.0 |

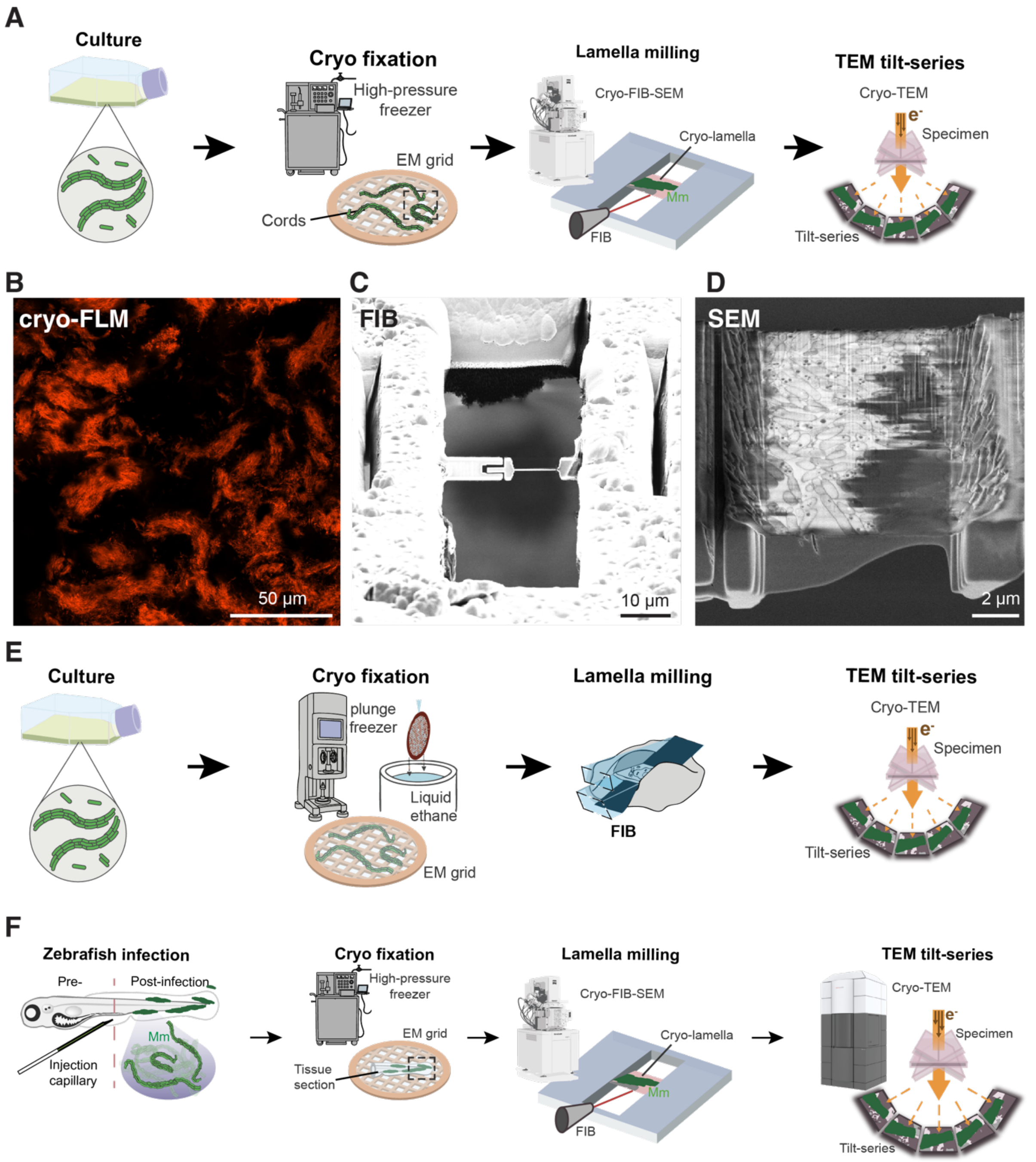


Fig. S1. Cryo-ET of mycobacterial cords. (A) Cryo-ET workflow for *M. tuberculosis* cords vitrified on electron microscopy grids by high-pressure freezing (HPF), thinned by focused ion beam milling and imaged by tomographic tilt-series data collection. (B-D) Correlative light and electron microscopy, using (B) cryo-fluorescence microscopy (cryo-FLM), (C) cryo-focused ion beam (cryo-FIB) milling and (D) cryo-scanning electron microscopy (cryo-SEM). (E) Cryo-ET workflow for *M. marinum* cords vitrified on electron microscopy grids by plunge freezing, thinned by focused ion beam milling and imaged by tomographic tilt-series data collection. (F) Cryo-ET workflow for *M. marinum* cords within infected zebrafish tissue, micro-sectioned onto grids, vitrified by HPF, thinned by focused ion beam milling and imaged by tomographic tilt-series data collection.

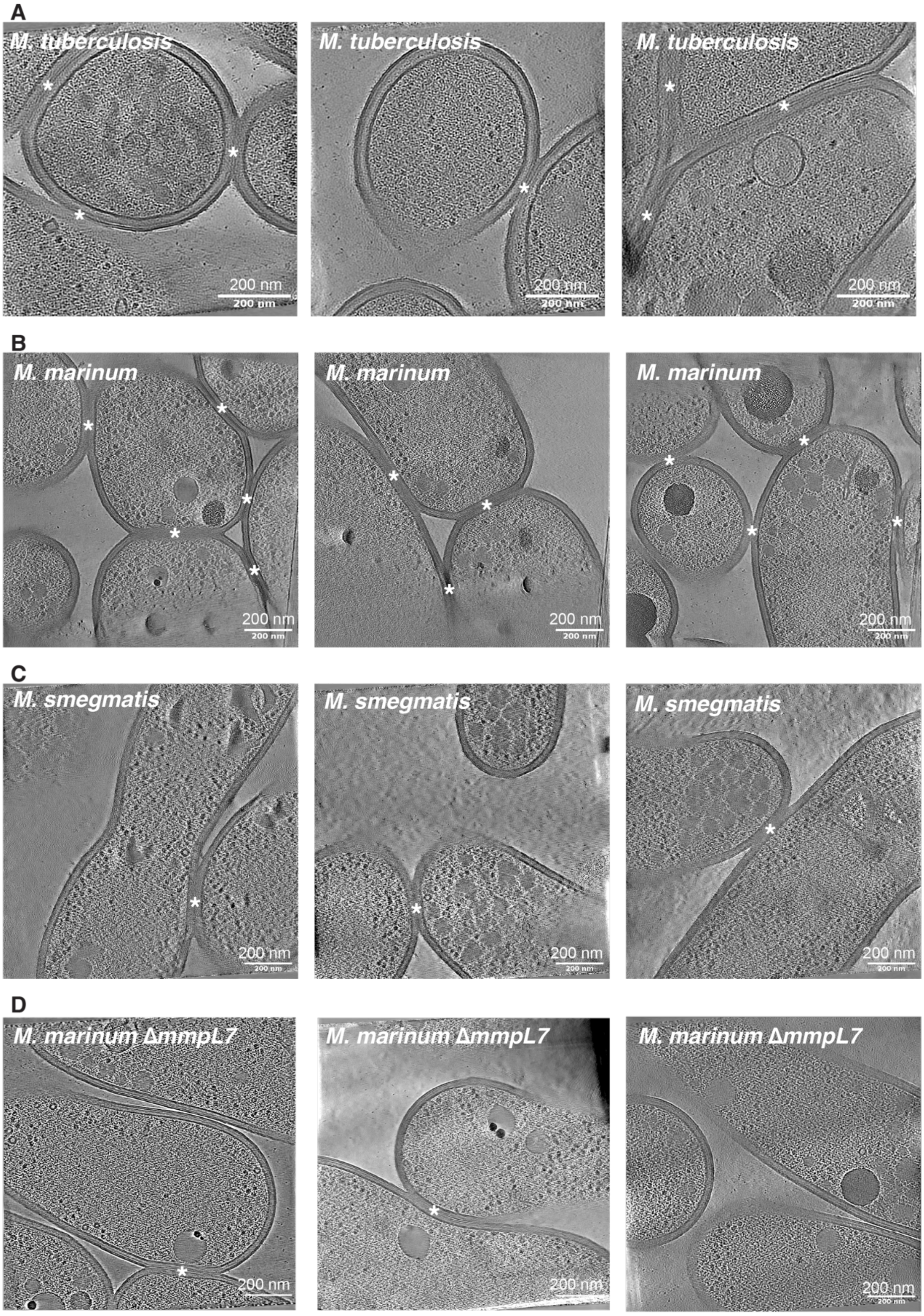


Fig. S2. Cryo-ET of wild-type and mutants of *M. tuberculosis*, *M. marinum* and *M. smegmatis*. (A) Cryo-ET of *M. tuberculosis* cords. Tomographic slices through cord cell-cell contacts are shown. (B) Cryo-ET of *M. marinum* cords. Tomographic slices through cord cell-cell contacts are shown. (C) Cryo-ET of *M. smegmatis* clumps. Tomographic slices through clumped cell-cell contacts are shown. (D) Cryo-ET of *M. marinum* Δ*mmpL7* clumps. Tomographic slices through clump cell-cell contacts are shown.

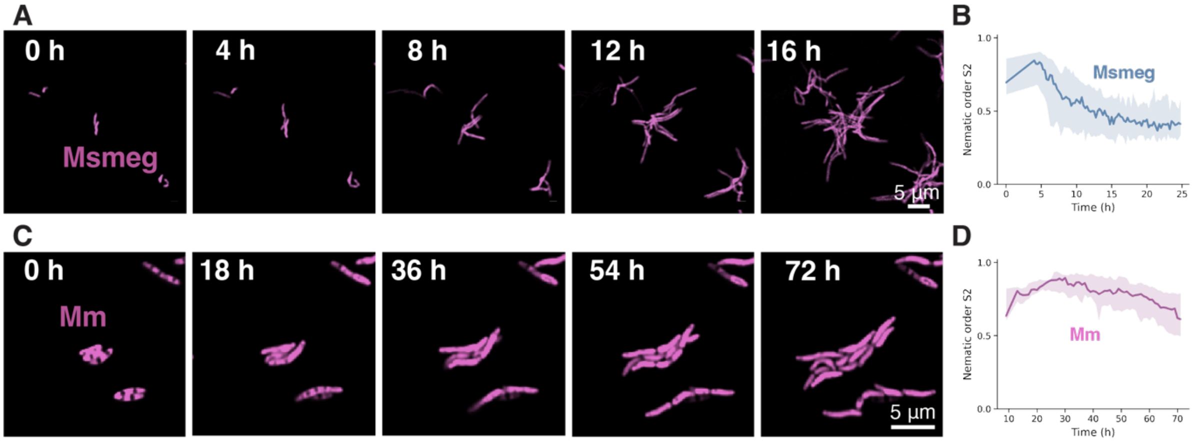


Fig. S3. Live cell confocal imaging of *M. smegmatis* and *M. marinum*. (A) Time-lapse fluorescence microscopy of *M. smegmatis* over 16 h, imaged every 15 minutes. (B) Local nematic order (S2) of *M. smegmatis* microcolonies over time. One time-lapse field per strain (85 frames at 15-minute intervals). n= 2,679 colony frames for colonies of 5–16 estimated cells. (C) Time-lapse fluorescence microscopy of *M. marinum* over 72 h, imaged every hour. (D) Local nematic order (S2) of *M. marinum* microcolonies over time. One time-lapse field per strain (72 frames at 60-minute intervals). 329 colony frames for colonies of 5–16 estimated cells.

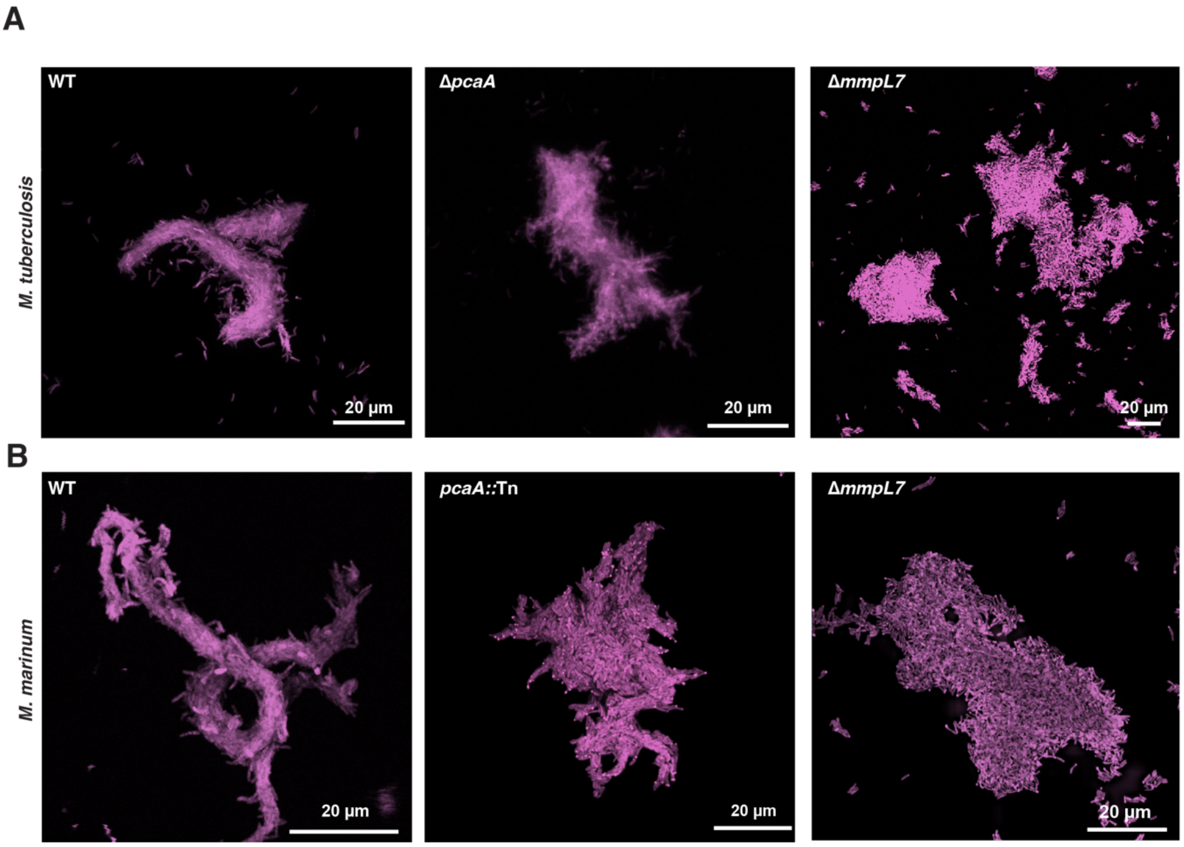


Fig. S4. Confocal imaging of wild-type and mutant *M. tuberculosis* and *M. marinum* colonies. (A-B) Additional fluorescence images of (A) *M. tuberculosis* colonies in wild-type, Δ*pcaA* and Δ*mmpL7* genetic backgrounds and (B) of *M. marinum* colonies in wild-type, *pcaA::*Tn and Δ*mmpL7* genetic backgrounds, using the same experimental conditions as Fig. 4A-B.

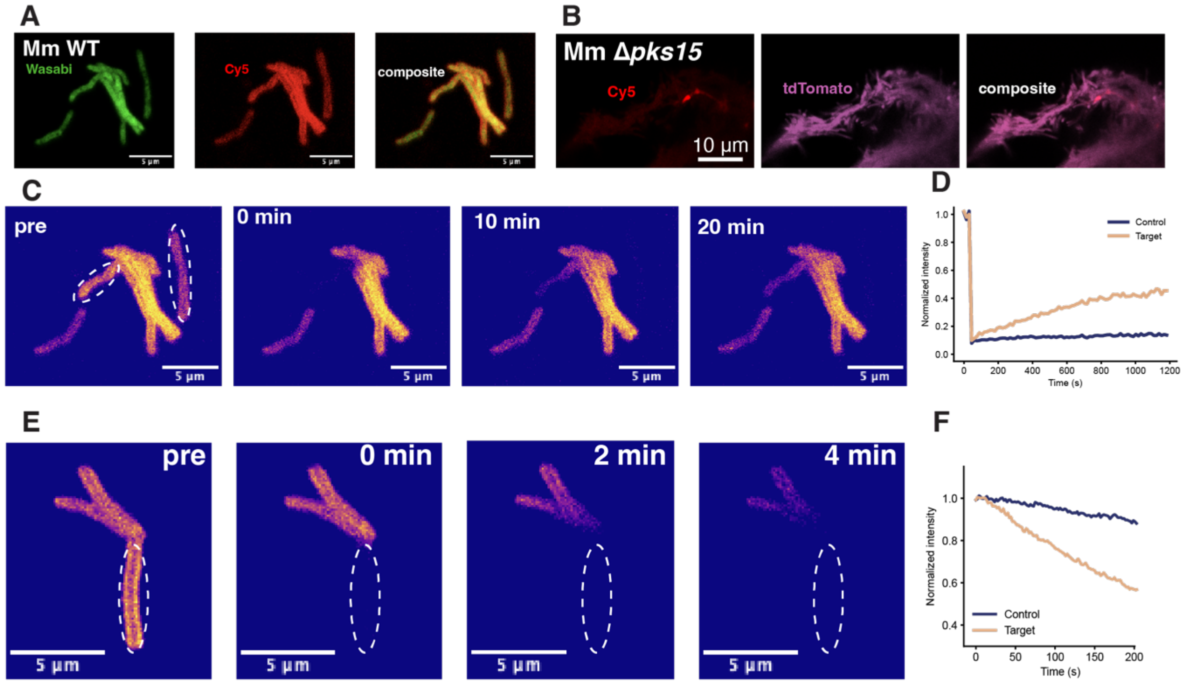


Fig. S5. Phenolic glycolipid diffuses through the shared mycomembrane compartment. (A) Early wild-type *M. marinum* cord (Wasabi, green, left), labelled with Cy5-PGL (red, middle) and the composite image shown (right). (B) *M. marinum* Δ*pks15* cords labelled with Cy5-PGL (red, left), cells shown (tdTomato, magenta, middle) and the composite image shown (right). (C-D) Additional representative fluorescence recovery after photobleaching (FRAP) experiment supporting Fig. 5, using the same experimental conditions. (C) A region within a cord (target) and a separate, non-contacting cell (control) were bleached and imaged over 12 minutes. Dashed outlines depict bleached regions. (D) Normalized fluorescence recovery for target and control regions. A single representative shown; recovery was observed in all cords tested (n=13). (E-F) Fluorescence loss in photobleaching (FLIP). (E) Continuous bleaching of one region of a cord. Scale bars, 5 μm. (F) Normalized intensity during continuous bleaching. A single representative experiment is shown; fluorescence loss was observed in all cords tested (n=4).

Movie S1. Movie showing sequential "Z"-slices through a reconstructed tomogram of a *M. tuberculosis* cord. Corresponds to static panel in Fig. 1E.

Movie S2. Movie showing sequential "Z"-slices through a reconstructed tomogram of a *M. marinum* cord. Representative tomogram slice is depicted in Fig. 2B.

Movie S3. Movie showing sequential "Z"-slices through a reconstructed tomogram of zebrafish tissue infected with *M. marinum*. Corresponds to panel Fig. 2G.

Movie S4. Movie showing sequential "Z"-slices through a reconstructed tomogram of a *M. smegmatis* clump. Corresponds to static panel in Fig. 3B.

Movie S5. Movie showing sequential "Z"-slices through a reconstructed tomogram of a *C. glutamicum* clump. Tomographic series corresponds to Fig. 3G.

Movie S6. Time-lapse imaging of *M. smegmatis*. This representative video corresponds to the images shown in Fig. 4A.

Movie S7. Live-cell imaging of *M. tuberculosis* capturing cell-cell rearrangement via sliding. Corresponds to the images shown in Fig. 4B.

Movie S8. Live-cell imaging of *M. tuberculosis* showing post-division cell-cell rearrangement via flipping. Corresponds to the images shown in Fig. 4B.

Movie S9. Live imaging of cellular rearrangements of *M. marinum,* with post-division cellular rearrangement via sliding. Corresponds to the images shown in Fig. 4C.

Movie S10. Live imaging of cellular rearrangements of *M. marinum*, capturing cell-cell rearrangement via flipping. Corresponds to the images shown in Fig. 4C.

Movie S11. Movie showing sequential "Z"-slices through a reconstructed tomogram of a mutant *M. marinum ΔmmpL7* clump. Corresponds to the single tomographic slice depicted in Fig. 5F.

### SI References


1. K. C. Murphy, K. Papavinasasundaram, C. M. Sassetti, Mycobacterial recombineering. *Methods Mol Biol* **1285**, 177–199 (2015).

2. M. I. Hayes, *et al.*, Mycobacteria trehalose dimycolate interactions with host Mincle remodel blood-brain barrier junctions for brain invasion. *Cell Rep* **44**, 116661 (2025).

3. K. Kelley, *et al.*, Waffle Method: A general and flexible approach for improving throughput in FIB-milling. *Nat Commun* **13**, 1857 (2022).

4. M. G. F. Last, L. Abendstein, L. M. Voortman, T. H. Sharp, Streamlining segmentation of cryo-electron tomography datasets with Ais. *Elife* **13**, RP98552 (2024).

5. Y.-T. Liu, *et al.*, IsoNet2 determines cellular structures at submolecular resolution without averaging. [Preprint] (2025).

6. M. G. F. So-Last, A. Burt, T. Hale, M. Allegretti, Easymode: general pretrained networks for cellular cryo-ET enable flexible approaches to subtomogram averaging. [Preprint] (2026). Available at: http://biorxiv.org/lookup/doi/10.64898/2026.05.19.726344 [Accessed 18 October 2026].

7. J. R. Kremer, D. N. Mastronarde, J. R. McIntosh, Computer visualization of three-dimensional image data using IMOD. *J Struct Biol* **116**, 71–76 (1996).

8. J. Schindelin, *et al.*, Fiji: an open-source platform for biological-image analysis. *Nat Methods* **9**, 676–682 (2012).

9. E. F. Pettersen, *et al.*, UCSF ChimeraX: Structure visualization for researchers, educators, and developers. *Protein Sci* **30**, 70–82 (2021).

10. U. H. Ermel, S. M. Arghittu, A. S. Frangakis, ArtiaX: An electron tomography toolbox for the interactive handling of sub-tomograms in UCSF ChimeraX. *Protein Sci* **31**, e4472 (2022).

11. B. Isbilir, A. Yeates, V. Alva, T. A. M. Bharat, Mapping the ultrastructural topology of the corynebacterial cell surface. *PLoS Biol* **23**, e3003130 (2025).

12. C. Paganelli, *et al.*, Scale Invariant Feature Transform as feature tracking method in 4D imaging: a feasibility study. *Annu Int Conf IEEE Eng Med Biol Soc* **2012**, 6543–6546 (2012).

13. S. van der Walt, *et al.*, scikit-image: image processing in Python. *PeerJ* **2**, e453 (2014).
